# Cancer PRSweb – an Online Repository with Polygenic Risk Scores (PRS) for Major Cancer Traits and Their Phenome-wide Exploration in Two Independent Biobanks

**DOI:** 10.1101/2020.01.22.915751

**Authors:** Lars G. Fritsche, Snehal Patil, Lauren J. Beesley, Peter VandeHaar, Maxwell Salvatore, Robert B. Peng, Daniel Taliun, Xiang Zhou, Bhramar Mukherjee

**Author notes:** Corresponding authors (L.G.F.), (B.M.).

## Abstract

To facilitate scientific collaboration on polygenic risk scores (PRS) research, we created an extensive PRS online repository for 49 common cancer traits integrating freely available genome-wide association studies (GWAS) summary statistics from three sources: published GWAS, the NHGRI-EBI GWAS Catalog, and UK Biobank-based GWAS. Our framework condenses these summary statistics into PRS using various approaches such as linkage disequilibrium pruning / p-value thresholding (fixed or data-adaptively optimized thresholds) and penalized, genome-wide effect size weighting. We evaluated the PRS in two biobanks: the Michigan Genomics Initiative (MGI), a longitudinal biorepository effort at Michigan Medicine, and the population-based UK Biobank (UKB). For each PRS construct, we provide measures on predictive performance, calibration, and discrimination. Besides PRS evaluation, the Cancer-PRSweb platform features construct downloads and phenome-wide PRS association study results (PRS-PheWAS) for predictive PRS. We expect this integrated platform to accelerate PRS-related cancer research.

## Introduction

Since 2005, genome-wide association studies (GWAS) have successfully uncovered many common genetic variants associated with a plethora of complex traits and disorders [1–3]. Translation of these findings into clinical practice to improve pre-symptomatic screening and patient care is a major aspiration in the research community. However, genetic risk factors for complex diseases like cancer usually have relatively small risk effects and/or low frequencies and thus only have limited ability as individual predictors of risk in the overall population. Alternatively, the integration of all common risk variants into a single biomarker, called a polygenic risk score (PRS), represents a widely used approach for potentially identifying high-risk individuals at the highest levels of PRS [4–6]. For example, it was shown that PRS for five common complex diseases (coronary artery disease, atrial fibrillation, type 2 diabetes, inflammatory bowel disease, and breast cancer) have the potential to detect individuals at significantly higher genetic risk [4] that might benefit from intensified screening efforts, prophylactic prevention or earlier treatment. Several challenges have to be overcome for constructing a PRS that incorporates state of the art scientific knowledge: one needs (1) summary statistics from an independent discovery GWAS with phenotype and ancestry matching the target study [7]; (2) individual-level genetic data of a sufficiently large cohort to correct for linkage disequilibrium (LD) between genetic variants; and (3) a computationally efficient method to calculate each PRS and to find the best PRS construct for the target cohort.

The gold standards for GWAS to define PRS constructs are independent, large GWAS analyses or GWAS meta-analyses. Full summary statistics enable exploration of the complete spectrum of PRS construction methods, e.g., those that determine the optimal inclusion p-value threshold of risk variants for prediction, which often deviates from the standard threshold for genome-wide significance (P-value ≤ 5×10^-8^). So far, several cancer GWAS research groups and consortia have openly shared their full GWAS summary statistics with the research community: ovarian carcinoma [8, 9], breast cancer [10, 11], prostate cancer [12], colorectal cancer [13], and cervical carcinoma [14]. Other groups have released variants that reached an arbitrarily chosen p-value threshold below genome-wide significance (e.g., P-value < 10^-5^)[15]. In addition to complete or partial GWAS summary statistics, lists of genome-wide significant hits are available for nearly all published GWAS results. The NHGRI-EBI GWAS Catalog [2] (https://www.ebi.ac.uk/gwas/) curates and stores published risk variants for a plethora of traits in a structured database, offering a convenient and efficient way to extract GWAS hits for automated processing.

Alternative and growing sources for publicly available GWAS summary statistics across a large ensemble of diseases use UK Biobank genotype and phenotype data [16], adjusting for population stratification and/or relatedness between individuals ([17]; http://www.nealelab.is/uk-biobank and https://www.leelabsg.org/resources). These biobank-based approaches accessed thousands of phenotypes and traits that were defined in efficient automated fashion, e.g., by ICD10 diagnosis category, with specific phenotype defining algorithms like PHESANT [18] or PHEWAS CODES [19], or even with consortium-based curated phenotype constructs using the content of the electronic health records (EHR) (FINNGEN; https://www.finngen.fi/en/researchers/clinical-endpoints).

Another important aspect of finding a suitable set of GWAS summary statistics for a PRS is the mapping of the discovery GWAS trait, here the cancer phenotype, with the trait of interest in the target study. GWAS efforts usually balance specificity and sample size to maximize power for discovery. Consequently, the analyzed phenotype definition might not necessarily represent an ideal match to the phenotype of the target study. Also, differences in diagnosis coding practices in EHR systems, e.g., the preference for certain diagnoses due to billing purposes, might limit the transferability of phenotype definitions across cohorts, even if the same coding systems were used [20].

The simplest form of PRS construction requires two things: a selected set of independent risk variants with estimated or weighted risk effect sizes (say 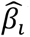), and genotype data of individuals genotyped at the selected sites (say *G_i_* where *i* ∈ a list *L*). A PRS can then be calculated for each individual as the sum of the weighted risk increasing alleles, namely 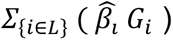.

PRS construction methods and their underlying variant selection procedures can roughly be categorized into four groups: (i) fixed P-value thresholds of independent risk variants, e.g., “GWAS hits,” variants that reached genome-wide significance (with *p* < 5 × 10^-8^; (ii) LD pruning (actually clumping) / P-value thresholding (“P&T”) of summary statistics that increases power by determining the most predictive P-value cut-off that can be above or below genome-significance [6]; (iii) genome-wide PRS that consider the full GWAS summary statistics after modeling LD, applying shrinkage or Bayesian approaches, e.g., LDpred and lassosum, [21–24] and (iv) methods that use individual-level data from a GWAS to determine an optimal set of independent predictors through Bayesian spike and slab or mixture priors [25]. The first two approaches typically use the originally reported effect sizes for weighting, while the latter two approaches model LD and/or shrink effect sizes. All methods require a reference panel for LD estimation that ideally resembles or matches the genotype data underlying the discovery GWAS source. Since most only have summary statistics and not individual-level data of the discovery study, we will use only the first three approaches for PRS construction, i.e., fixed P-value thresholds, LD pruning / P-value thresholding, and Lassosum.

PRS have increasingly been used in cancer risk prediction and stratification. A brief survey of PRS related literature in PubMed shows that ∼15% of all PRS articles are related to cancer, with 67% of cancer PRS papers focusing on common cancers (defined by the US National Cancer Institute (NCI; https://www.cancer.gov/types/common-cancers; estimated incidence of 40,000 or more in the United States in 2019). As of November 9, 173 PubMed articles on PRS and cancer have been published in 2019, more than double the previous high of 86 set in 2018, indicating the rapid growth in collection, curation, and generation of genetic data. These studies typically employ construction methods (i) and (ii) as described above, although joint variant models are becoming more common because they generally outperform methods (i) and (ii) and advanced software has made joint modeling more computationally efficient for large sample sizes [26, 27]. Several publications constructed PRS for cancer traits using different methods [28–30] and described their PRS methodology. However, very few share the variants selected and their corresponding weights, making it a challenge to compare or replicate PRS results in different cohorts. The Polygenic Score Catalog (http://www.pgscatalog.org) is a resource under active development to help researchers share, apply, and evaluate PRS. However, this resource primarily relies on external PRS sources and currently considers only 21 traits (including only four cancer traits: ovarian, colorectal, prostate and breast cancer), and no validation is carried out in large biobanks.

The primary goal of this study is the generation of PRS constructs for common groupings of cancer by using published, freely available cancer GWAS summary statistics and established PRS methods and genetic data from two large biobanks: the Michigan Genomics Initiative (MGI) and the UK Biobank (UKB) (**Table 1**). We explore hundreds of PRS constructs and offer optimized predictive PRS (in terms of maximal increase in an *R^2^*-type metric) for 49 cancers. The resulting repository of cancer PRS is made available online via an interactive platform, called *Cancer PRSweb* (http://prsweb.sph.umich.edu). In this platform, we accompany each GWAS source / PRS method combination with its downloadable constructs and performance metrics (like area under the receiver operating curve, tail enrichment, and Brier score), and we offer insights into secondary trait associations through screening of hundreds of cancer and non-cancer phenotypes of the EHR-derived phenomes of MGI and UKB (**Table 1**). We also make the summary statistics for the phenome-wide association study (PheWAS) available. Thus, this centralized and unified platform is a timely attempt to accelerate cancer research related to PRS.

**Table 1.** Demographics and clinical characteristics of the analytic datasets. The provided characteristics are based on the European (MGI) and White British (UKB) subjects for which phenotype and imputed genotype data were available.

| Characteristic | MGI | UKB |
| --- | --- | --- |
| Total participants | 38,360 | 408,595 |
| Females, n (%) | 20,141 (52.5%) | 220,896 (54.1%) |
| Mean age, years (S.D.) | 56.8 (16.2) | 56.9 (8.0) |
| Median number of visits per participant | 45 | not available |
| Median time (years) between first and last visit | 5.5 | not available |
| Median number of unique ICD9 codes | 36* | 2 |
| Median number of unique ICD10 codes | 31* | 6 |
| Number of Phecodes with more than 50 cases | 1,689 | 1,419 |
| Any cancer diagnosis | 20,751 (54.1%) | 69,190 (16.9%) |
| *20 Most Common Cancer Traits in MGI (Phecode)* |  |  |
| Basal cell carcinoma (172.21) | 2,988 (7.79%) | not available |
| Melanomas of skin, dx or hx (172.1) | 2,701 (7.04%) | 2,682 (0.66%) |
| Breast cancer [female] (174.1) | 2,605 (12.93%) | 12,483 (5.65%) |
| Cancer of prostate (185) | 2,432 (13.35%) | 5,977 (3.18%) |
| Squamous cell carcinoma (172.22) | 1,917 (5.00%) | not available |
| Cancer of bladder (189.2) | 1,575 (4.11%) | 2,413 (0.59%) |
| Colorectal cancer (153) | 1,196 (3.12%) | 4,585 (1.12%) |
| Non-Hodgkins lymphoma (202.2) | 1,141 (2.97%) | 1,810 (0.44%) |
| Malignant neoplasm of kidney, except pelvis (189.11) | 1,083 (2.82%) | 1,033 (0.25%) |
| Colon cancer (153.2) | 941 (2.45%) | 3,108 (0.76%) |
| Myeloproliferative disease (200) | 886 (2.31%) | 992 (0.24%) |
| Cancer of bronchus; lung (165.1) | 874 (2.28%) | 2,232 (0.55%) |
| Thyroid cancer (193) | 798 (2.08%) | 347 (0.08%) |
| Malignant neoplasm of rectum, rectosigmoid junction, and anus (153.3) | 669 (1.74%) | 2,167 (0.53%) |
| Malignant neoplasm of uterus (182) | 643 (3.19%) | 1,285 (0.58%) |
| Nodular lymphoma (202.21) | 632 (1.65%) | 365 (0.09%) |
| Cancer of tongue (145.2) | 550 (1.43%) | 310 (0.08%) |
| Leukemia (204) | 545 (1.42%) | 1,665 (0.41%) |
| Cancer of brain (191.11) | 483 (1.26%) | 525 (0.13%) |
| Cervical cancer (180.1) | 430 (1.12%) | 272 (0.12%) |

\* ICD9/10-CM codes S.D. standard deviation

Our repository contributes to the new and necessary work of democratizing PRS constructions and applications for several cancers under a uniform analytic framework to eventually develop transferable risk scores with clinical utility. We also offer phenome-wide exploration of PRS association through PRS-PheWAS, a tool previously introduced by this group [31, 32].

## Results

### PRS Construction

We screened the GWAS Catalog, PubMed, and UK Biobank GWAS efforts for any cancer GWAS summary statistics that were reported for European ancestry, to match the predominantly European cohorts of MGI and UKB, and that were openly available, i.e., did not require contacting the main authors or any form of written approval process. We identified 232 source sets that reported complete information for each tested single nucleotide polymorphisms (SNP) (position [and/or dbSNP ID], effect allele, effect estimate, p-value, and, ideally, effect allele frequency). We obtained 188 SNP sets based on UKB GWAS, 29 based on excerpts from the GWAS Catalog, and 20 from large GWAS or GWAS meta-analyses (**Table S1 & S2**).

We manually matched the traits of the identified cancer GWAS to cancer traits of MGI and UKB PheWAS-codes and analyzed each GWAS source separately, generating PRS for each. The discovery GWAS traits of the 232 source sets approximated 68 cancer PheWAS-codes of the MGI phenome and 21 PheWAS-codes in the UKB phenome (**Table S1 & S2**). Following the scheme in **Figure 1** and **Table 2**, we generated PRS using the “P & T” and/or “lassosum” approach and also generated PRS using fixed P-value thresholds after LD clumping (P-value ≤ 5×10^-5^, 5×10^-6^, 5×10^-7^, 5×10^-8^ [“GWAS Hits”], or 5×10^-9^). Using these methods and the available GWAS sources, we generated a total of 1,292 PRS (1,080 PRS for the MGI cohort and 212 PRS for the UKB cohort) (**Table S3**).

**Figure 1.**
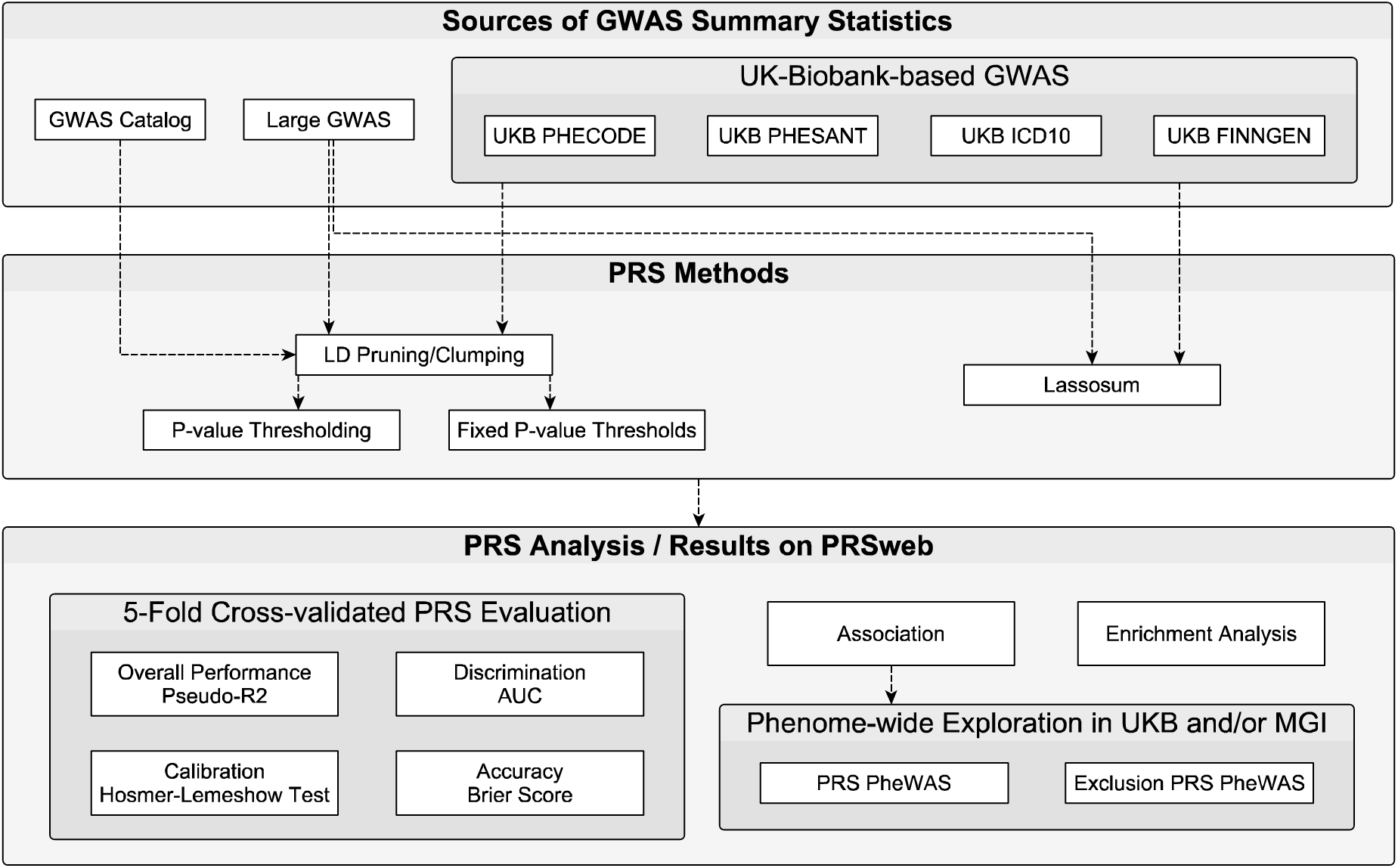
Schematic overview of PRS generation and analysis.

**Table 2.**
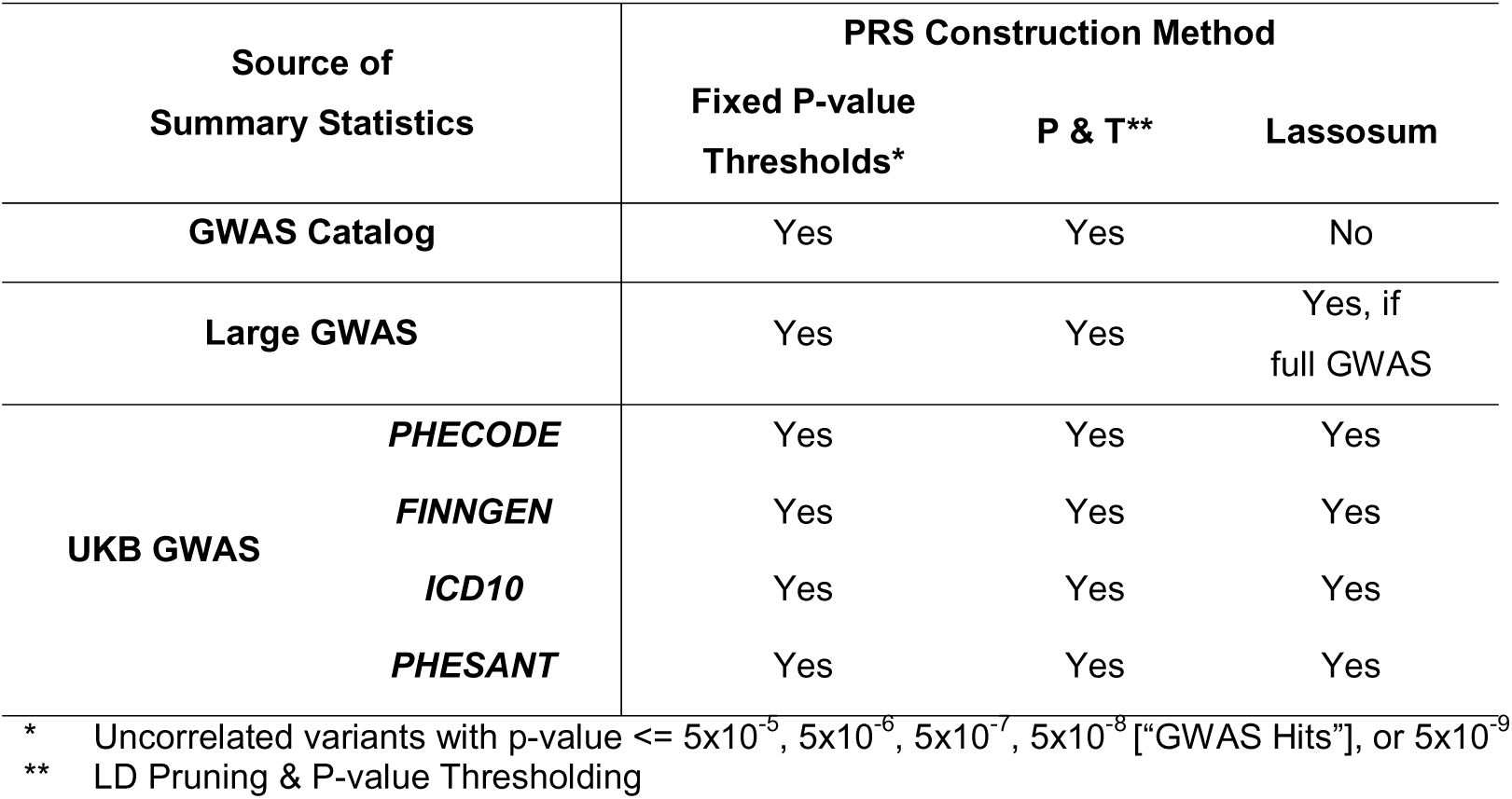
Overview of GWAS sources and PRS construction methods. Multiple PRS were constructed per trait of interest depending on availability of GWAS summary statistics.

### PRS Evaluation

We tested the association between each PRS and its corresponding cancer trait and evaluated each PRS in terms of performance (pseudo-R^2^), accuracy (Brier score), discrimination (area under the receiver operating characteristic curve [AUC]), and calibration (Hosmer-Lemeshow test). Finally, we tested their utility for risk stratification, i.e., their ability to enrich cases in five selected top percentiles (1, 2, 5, 10, and 25%) versus the rest of the PRS distribution (**Figure 1**).

As an initial filtering step, we removed 625 PRS (48% of total PRS considered) that were not significantly (603 PRS with P > 0.05) or negatively (241 PRS) associated with their corresponding cancer trait in MGI and/or UKB. The majority of these filtered PRS were either based on discovery GWAS with small sample sizes that often did not identify any genome-wide significant hits or were evaluated for diseases with few cases or both, indicating a potential lack of power in our analysis. A total of 667 PRS for 49 different cancer traits were positively and significantly associated with their corresponding cancers in MGI (478 PRS; 49 cancer traits) and UKB (189 PRS; 19 cancer traits) (**Table S3**).

#### Comparison of Performance Metrics

In general, we found that the ranking by pseudo-R^2^ ensured strong performance across other metrics related to discrimination, accuracy, and overall association of PRS constructs for their specific cancers. Conversely, the enrichment analyses in the extreme PRS percentiles (e.g., top 5% versus rest) was not always concordant with the selection of optimal PRS based on pseudo-R^2^, showing that performance in the extreme tails could be optimized by a modified criterion that focuses on extremes of the risk distribution [33].

An example evaluation is shown in **Table 3**. Here we compare PRS across seven construction methods (lassosum, P&T, and five fixed P-value thresholds) that were all based on a single summary statistics source, a large GWAS on overall breast cancer [11]. In MGI, we observed that the lassosum-based PRS (44,815 SNPs) had the best performance (highest pseudo-R^2^ = 0.057), the highest accuracy (Brier score = 0.137), the best discrimination between breast cancer cases and controls (AUC = 0.635 [95% confidence interval (CI): 0.624, 0.647]), and showed the strongest association with breast cancer itself (odds ratio [OR]_continuous PRS_ = 1.66 [95% CI: 1.58,1.73]). In this scenario, modeling LD information with lassosum retained more information than LD clumping [22], even though, unlike the other methods, lassosum only considered autosomal variants.

**Table 3.** Comparison of PRS methods on Breast Cancer PRS Performance in MGI (2,605 breast cancer cases and 12,548 controls). PRS are based on the BCAC Consortium GWAS on overall breast cancer [11]. Shaded cells indicate best performing PRS according to the corresponding metrics for MGI or UKB.

| Cohort | Method<br>Tuning Paramter | # SNPs | Pseudo-R2 | Hosmer-Lemeshow<br>P | Brier Score | AUC<br>(95% CI) | Odds Ratio<br>continuous PRS<br>(95% CI) | Odds Ratio<br>Top 1%<br>(95% CI) |
| --- | --- | --- | --- | --- | --- | --- | --- | --- |
| MGI | Lassosum<br>$s = 0.5, \lambda = 0.0055$ | 44,815 | 0.057 | 0.013 | 0.137 | 0.635<br>(0.624,0.647) | 1.66<br>(1.58,1.73) | 3.38<br>(2.42,4.71) |
| | P&T<br>$P \leq 0.00032$ | 2,723 | 0.046 | 0.018 | 0.138 | 0.626<br>(0.615,0.638) | 1.59<br>(1.52,1.67) | 3.58<br>(2.56,4.96) |
| | Fixed Threshold<br>$P \leq 5e-05$ | 1,307 | 0.045 | 0.008 | 0.138 | 0.625<br>(0.613,0.637) | 1.58<br>(1.51,1.65) | 3.41<br>(2.43,4.73) |
| | Fixed Threshold<br>$P \leq 5e-06$ | 712 | 0.044 | 0.16 | 0.138 | 0.622<br>(0.61,0.634) | 1.56<br>(1.49,1.63) | 3.35<br>(2.4,4.66) |
| | Fixed Threshold<br>$P \leq 5e-07$ | 464 | 0.043 | 0.18 | 0.139 | 0.621<br>(0.609,0.633) | 1.55<br>(1.49,1.63) | 3.69<br>(2.65,5.12) |
| | Fixed Threshold<br>$P \leq 5e-08$ | 334 | 0.041 | 0.084 | 0.139 | 0.619<br>(0.608,0.631) | 1.54<br>(1.48,1.61) | 3.77<br>(2.71,5.23) |
| | Fixed Threshold<br>$P \leq 5e-09$ | 264 | 0.04 | 0.02 | 0.139 | 0.618<br>(0.606,0.629) | 1.53<br>(1.47,1.6) | 3.15<br>(2.25,4.38) |
| UKB | Lassosum<br>$s = 0.9, \lambda = 0.0043$ | 286,144 | 0.047 | 3.50E-27 | 0.0808 | 0.645<br>(0.64,0.65) | 1.71<br>(1.68,1.74) | 4.48<br>(3.98,5.03) |
| | P&T<br>$P \leq 1e-04$ | 1,682 | 0.04 | 3.10E-23 | 0.0811 | 0.635<br>(0.63,0.64) | 1.64<br>(1.61,1.67) | 3.57<br>(3.16,4.03) |
| | Fixed Threshold<br>$P \leq 5e-06$ | 712 | 0.04 | 2.40E-20 | 0.0811 | 0.633<br>(0.628,0.638) | 1.62<br>(1.59,1.66) | 3.8<br>(3.37,4.28) |
| | Fixed Threshold<br>$P \leq 5e-05$ | 1,307 | 0.04 | 2.70E-18 | 0.0811 | 0.633<br>(0.628,0.638) | 1.63<br>(1.6,1.66) | 3.63<br>(3.21,4.09) |
| | Fixed Threshold<br>$P \leq 5e-07$ | 464 | 0.039 | 3.80E-20 | 0.0812 | 0.632<br>(0.627,0.637) | 1.61<br>(1.58,1.64) | 3.9<br>(3.45,4.39) |
| | Fixed Threshold<br>$P \leq 5e-08$ | 334 | 0.037 | 1.00E-18 | 0.0813 | 0.629<br>(0.624,0.634) | 1.59<br>(1.56,1.62) | 3.71<br>(3.28,4.18) |
| | Fixed Threshold<br>$P \leq 5e-09$ | 264 | 0.035 | 6.60E-15 | 0.0813 | 0.626<br>(0.621,0.631) | 1.57<br>(1.54,1.6) | 3.42<br>(3.02,3.87) |

The enrichment of cases in the top 1% compared to the rest was more pronounced for the “GWAS hits” PRS (Fixed Threshold P <= 5e-08; 334 SNPs; OR_Top1%_ 3.77 [95% CI: 2.71,5.23]) than for the lassoum-PRS (OR_Top1%_ = 3.38 [95% CI: 2.42,4.71]). We found signs that the logistic regression model that we used for the evaluation might be misspecified for some PRS, e.g., four of the seven PRS were not well-calibrated according to the Goodness-of-Fit test statistics (Hosmer-Lemeshow P <= 0.05) (**Table 3**). In UKB, we observed an identical ranking of PRS methods but we noted several differences with MGI. First, the tuning parameters of the lassosum-PRS and the P&T-PRS differed between MGI and UKB, resulting in a different number of included variants (lassosum: MGI: 44,815 variants versus UKB: 286,144 variants; P&T: MGI 2,723 variants versus UKB: 1,682 variants) (**Table 3**). Closer inspection of the underlying tuning parameter optimization revealed comparable parameter ranking for lassosum and P&T, suggesting that optimizations seem cohort-specific but stable, i.e., a predictive PRS established in UKB might perform similarly well in MGI and vice versa (Spearman’s rank correlation > 0.98) (**Figure S1**).

#### Comparison across GWAS Sources

We also explored the influence of various GWAS sources on the predictive performance of PRS. As an illustrative example, we again focus on breast cancer PRS, but now consider PRS constructed from different breast cancer GWAS sources, using for each source the method that yielded the highest pseudo-R^2^ (**Table 4**). In MGI, the PRS (lassosum) of the largest available GWAS (122,977 cases and 105,974 controls) yielded the best performance across most PRS metrics (e.g. pseudo-R^2^ = 0.057, AUC = 0.635 [0.624,0.647]). The GWAS Catalog-PRS (P&T), which included 79 top hits from 18 different GWAS [11, 34–51], was ranked second (pseudo-R^2^ = 0.034) and showed significantly inferior discrimination ability (AUC 0.603 [0.591,0.615]). The case enrichment in the top 1% was pronounced but not significantly different from the top-ranked PRS (OR_Top1%_[GWAS Catalog] = 3.38 [2.42,4.71] versus OR_Top1%_[Large GWAS] = 3.52 [2.52,4.89]). The four UKB GWAS-based PRS (all based on lassosum) followed next and showed similar performances (pseudo-R^2^: 0.029 – 0.022; AUC between 0.603 – 0.586 with overlapping confidence intervals) and could be ranked according to their effective sample sizes. Most interestingly, the “UKB PheCode” PRS (6,977 variants) could differentiate cases and controls as well as the GWAS Catalog PRS, which was based on 79 independent risk variants with P <= 2.5e-08 reported in 18 GWAS (both AUC 0.603 [0.591,0.615]). This suggested that biobank-based PRS can be a viable alternative for PRS construction, especially if summary statistics from a large disease-specific GWAS are unavailable (**Table 4**). A detailed comparison of GWAS sources across the 49 cancer traits in MGI is available in **Table S3**.

**Table 4.** Influence of GWAS sources on Breast Cancer PRS Performance in MGI. Shaded cells indicate best performing PRS according to the corresponding metrics for MGI or UKB.

| Cohort | GWAS Source<br>(Effective Sample Size*) | Method<br>Tuning Parameter | # SNPs | Pseudo<br>-R<sup>2</sup> | Hosmer-Lemeshow<br>P | Brier<br>Score | AUC<br>(95% CI) | Odds Ratio<br>continuous<br>PRS | Odds<br>Ratio<br>Top 1% |
| --- | --- | --- | --- | --- | --- | --- | --- | --- | --- |
| MGI | Large GWAS<br>[11]<br>(113,844) | Lassosum<br>s = 0.5, $\lambda$ = 0.0055 | 44,815 | 0.057 | 0.013 | 0.137 | 0.635<br>(0.624,0.647) | 1.66<br>(1.58,1.73) | 3.38<br>(2.42,4.71) |
|  | GWAS Catalog<br>[11, 34-51]<br>(variable) | P&T<br>P <= 2.5e-08 | 79 | 0.034 | 0.35 | 0.139 | 0.603<br>(0.591,0.615) | 1.46<br>(1.39,1.52) | 3.52<br>(2.52,4.89) |
| | UKB GWAS<br>PheCode<br>(23,838) | Lassosum<br>s = 0.5, $\lambda$ = 0.014 | 6,977 | 0.029 | 0.012 | 0.140 | 0.603<br>(0.591,0.615) | 1.44<br>(1.38,1.50) | 2.28<br>(1.60,3.22) |
| | UKB GWAS<br>FINNGEN<br>(18,375) | Lassosum<br>s = 0.9, $\lambda$ = 0.014 | 31,252 | 0.028 | 0.63 | 0.140 | 0.599<br>(0.587,0.611) | 1.41<br>(1.35,1.47) | 2.23<br>(1.56,3.14) |
| | UKB GWAS<br>PHESANT<br>(15,282) | Lassosum<br>s = 0.5, $\lambda$ = 0.018 | 5,025 | 0.023 | 0.51 | 0.140 | 0.586<br>(0.574,0.598) | 1.36<br>(1.30,1.42) | 2.55<br>(1.79,3.57) |
| | UKB GWAS<br>ICD10<br>(15,792) | Lassosum<br>s = 0.9, $\lambda$ = 0.018 | 7,388 | 0.022 | 0.027 | 0.140 | 0.588<br>(0.576,0.600) | 1.35<br>(1.29,1.41) | 2.34<br>(1.65,3.28) |
| UKB | Large GWAS<br>[11]<br>(113,844) | Lassosum<br>s = 0.9, $\lambda$ = 0.0043 | 286,144 | 0.047 | 3.5E-27 | 0.0808 | 0.645<br>(0.640,0.65) | 1.71<br>(1.68,1.74) | 4.48<br>(3.98,5.03) |
|  | GWAS Catalog<br>[11, 34-51]<br>(variable) | P&T<br>P <= 2.5e-08 | 79 | 0.024 | 4.4E-07 | 0.0818 | 0.605<br>(0.600,0.610) | 1.46<br>(1.43,1.48) | 2.65<br>(2.32,3.02) |

\* Effective sample size: $2 / (1/\text{\#cases} + 1/\text{\#controls})$ ;

#### Comparison of Performance Across Methods

First, we explored the benefit of p-value thresholding for the pre-filtered risk variants of the GWAS Catalog. Compared to the GWAS hits only approach, i.e., only perform LD-clumping of risk variants with P <= 5×10^-8^, the P-value thresholding step of the P&T PRS construction improved PRS performance, as previously reported [52]. This implied that P-value thresholding might to be beneficial even for the relatively sparse sets of GWAS hits reported in the GWAS Catalog (**Figure S2).**

The P&T approach will, by definition, also cover fixed p-value thresholds in its tuning parameter optimization, i.e., the final P&T PRS will be based on the p-value threshold with the highest pseudo-R^2^ and thus perform at least as well as any tested fixed p-value thresholds. Therefore, we limited our next comparison of PRS methods for full summary statistics to P&T and lassosum PRS. We assessed both methods for different GWAS sources in MGI (58 PRS) and UKB (12 PRS). We found that both methods ranked comparably, i.e., a GWAS source that produced a lassosum PRS with high pseudo-R^2^ also produced a P&T PRS with high pseudo-R^2^ and vice versa (Spearman’s rank correlation: rho > 0.937; (**Figure S3**).

#### Comparison of Performance across Cancers

Next, we were interested in comparisons between PRS across traits to assess overall performance and general differences between cancer traits. **Table 5** shows the top-ranked PRS for the 20 most common cancer traits in MGI and highlights the different properties of the generated PRS. The PRS vary in their numbers of included SNPs and their abilities to distinguish cases from controls or to enrich cases in the top percentiles. The AUC of the presented PRS was highest for cancer of prostate PRS (AUC=0.664 [0.652, 0.676]) and lowest for the nodular lymphoma PRS (0.535 [95% CI:0.512, 0.559]). Significant enrichment of cases in the top 1% ranged from OR of 6.54 (95% CI: 4.41, 9.79; cancer of prostate) to 2.13 (95% CI: 1.03, 4.01; leukemia). Due to limited sample sizes in the top percentiles, we could not detect significant enrichment for most of the rarer cancers.

**Table 5:**
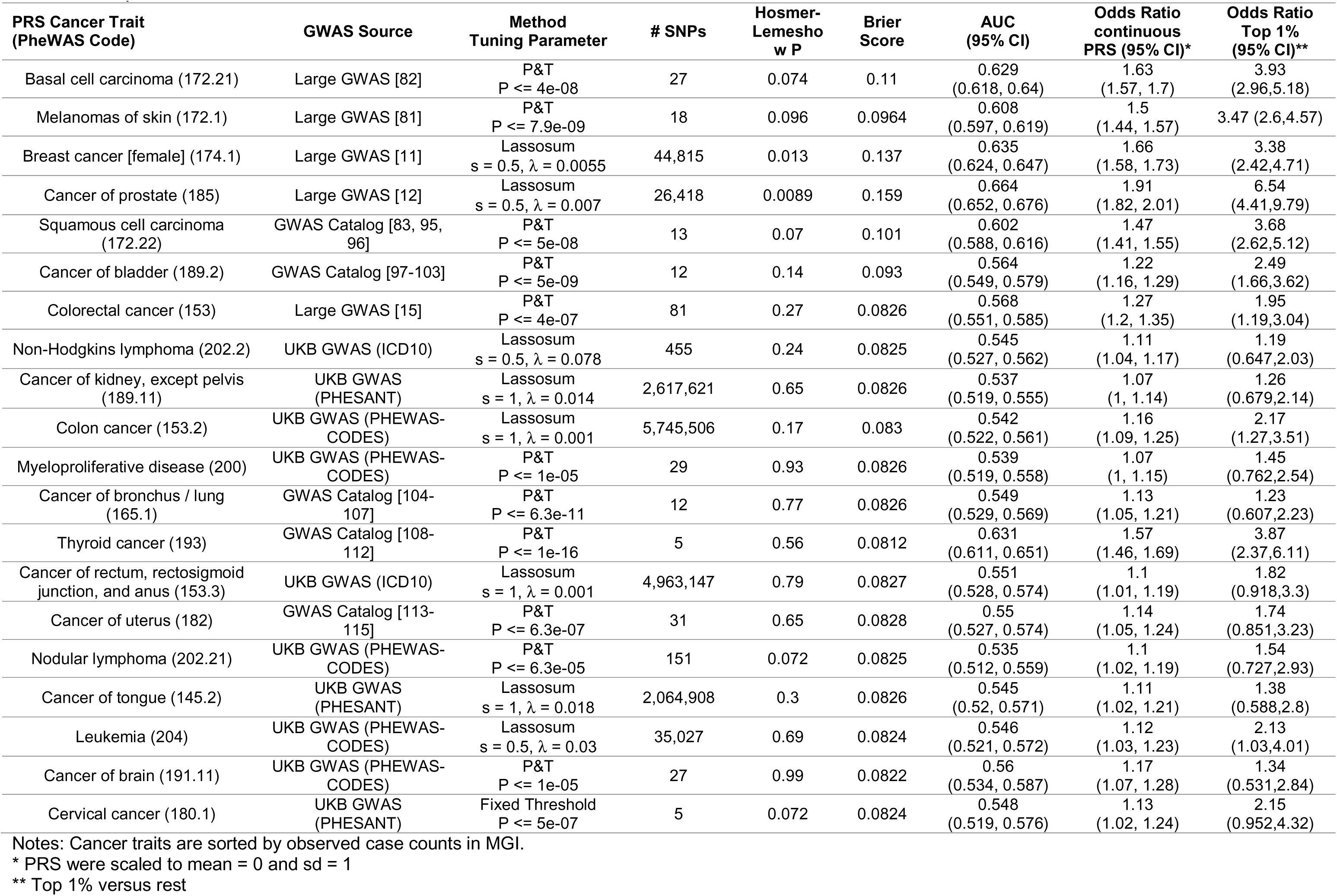
Top PRS for the 20 most common cancer traits in MGI (see Table 1)

**Table 6:** Best performing PRS for the 19 cancer traits in UKB

| PRS Cancer Trait<br>(PheWAS Code) | GWAS Source | Method<br>Tuning Parameter | # SNPs | Hosmer-Lemeshow<br>P | Brier<br>Score | AUC<br>(95% CI) | Odds Ratio<br>continuous PRS<br>(95% CI)* | Odds Ratio<br>Top 1%<br>(95% CI)** |
| --- | --- | --- | --- | --- | --- | --- | --- | --- |
| Breast cancer [female]<br>(174.1) | Large GWAS<br>[11] | Lassosum<br>s = 0.9, $\lambda$ = 0.0043 | 286,144 | 3.50E-27 | 0.0808 | 0.645<br>(0.64, 0.65) | 1.71<br>(1.68, 1.74) | 4.48<br>(3.98,5.03) |
| Cancer of prostate (185) | Large GWAS<br>[12] | Lassosum<br>s = 0.9, $\lambda$ = 0.0055 | 178,259 | 1.90E-33 | 0.08 | 0.698<br>(0.691, 0.705) | 2.1<br>(2.04, 2.16) | 6.28<br>(5.34,7.37) |
| Colorectal cancer (153) | Large GWAS<br>[15] | P&T<br>P <= 7.8e-06 | 87 | 0.0062 | 0.0813 | 0.618<br>(0.61, 0.627) | 1.55<br>(1.5, 1.59) | 3.48<br>(2.84,4.25) |
| Melanomas of skin (172.1) | GWAS Catalog<br>[81, 116-121] | P&T<br>P <= 1e-06 | 29 | 0.023 | 0.0815 | 0.611<br>(0.6, 0.623) | 1.5<br>(1.44, 1.56) | 2.94<br>(2.22,3.86) |
| Cancer of bladder (189.2) | GWAS Catalog<br>[97-103] | P&T<br>P <= 7e-07 | 15 | 0.2 | 0.082 | 0.577<br>(0.565, 0.589) | 1.32<br>(1.27, 1.38) | 2.24<br>(1.62,3.04) |
| Cancer of other lymphoid,<br>histiocytic tissue (202) | GWAS Catalog<br>[122, 123] | P&T<br>P <= 2.5e-06 | 6 | 0.76 | 0.0823 | 0.532<br>(0.519, 0.545) | 1.12<br>(1.07, 1.16) | 1.98<br>(1.41,2.72) |
| Cancer of bronchus / lung<br>(165.1) | GWAS Catalog<br>[104-107] | P&T<br>P <= 2.5e-08 | 19 | 0.47 | 0.0823 | 0.558<br>(0.545, 0.57) | 1.22<br>(1.17, 1.28) | 1.75<br>(1.21,2.46) |
| Non-Hodgkins lymphoma<br>(202.2) | GWAS Catalog<br>[124-128] | P&T<br>P <= 1e-09 | 10 | 0.21 | 0.082 | 0.558<br>(0.544, 0.573) | 1.23<br>(1.18, 1.29) | 1.96<br>(1.32,2.83) |
| Cancer of uterus (182) | GWAS Catalog<br>[113-115] | P&T<br>P <= 5e-08 | 18 | 0.55 | 0.0821 | 0.571<br>(0.554, 0.587) | 1.27<br>(1.2, 1.35) | 1.98<br>(1.24,3.04) |
| Cancer of kidney, except<br>pelvis (189.11) | GWAS Catalog<br>[129, 130] | P&T<br>P <= 5e-08 | 12 | 0.084 | 0.0826 | 0.54<br>(0.521, 0.558) | 1.11<br>(1.04, 1.17) | 1.45<br>(0.797,2.44) |
| Cancer of ovary (184.11) | Large GWAS [9] | Lassosum<br>s = 0.9, $\lambda$ = 0.0089 | 312,194 | 1 | 0.0821 | 0.573<br>(0.554, 0.592) | 1.28<br>(1.2, 1.37) | 2.3<br>(1.36,3.7) |
| Pancreatic cancer (157) | GWAS Catalog<br>[131-135] | P&T<br>P <= 7.9e-08 | 18 | 0.95 | 0.082 | 0.588<br>(0.565, 0.61) | 1.35<br>(1.25, 1.47) | 3.39<br>(1.94,5.66) |
| Cancer of brain and<br>nervous system (191.1) | GWAS Catalog<br>[136-142] | P&T<br>P <= 5e-07 | 25 | 0.15 | 0.0814 | 0.628<br>(0.604, 0.651) | 1.51<br>(1.4, 1.64) | 3.84<br>(2.14,6.58) |
| Multiple myeloma (204.4) | GWAS Catalog<br>[143-148] | P&T<br>P <= 6.3e-08 | 24 | 0.72 | 0.0817 | 0.597<br>(0.572, 0.621) | 1.41<br>(1.29, 1.54) | 2.26<br>(1.13,4.16) |
| Cancer of brain (191.11) | GWAS Catalog<br>[141, 142, 149] | P&T<br>P <= 6e-06 | 15 | 0.099 | 0.081 | 0.638<br>(0.612, 0.663) | 1.58<br>(1.45, 1.72) | 3.95<br>(2.16,6.9) |
| Lymphoid leukemia, chronic<br>(204.12) | GWAS Catalog<br>[150-156] | P&T<br>P <= 7.9e-08 | 33 | 0.0022 | 0.0794 | 0.69<br>(0.667, 0.714) | 1.98<br>(1.8, 2.17) | 5.82<br>(3.31,9.99) |
| Thyroid cancer (193) | GWAS Catalog<br>[108-112] | P&T<br>P <= 1e-16 | 5 | 0.27 | 0.0797 | 0.643<br>(0.612, 0.675) | 1.67<br>(1.5, 1.86) | 7.29<br>(3.79,13.7) |
| Cancer of testis (187.2) | GWAS Catalog<br>[157-164] | P&T<br>P <= 5e-06 | 44 | 0.00013 | 0.0805 | 0.677<br>(0.646, 0.708) | 1.87<br>(1.65, 2.12) | 2.98<br>(1.21,6.47) |
| Hodgkin's disease (201) | GWAS Catalog<br>[165-169] | P&T<br>P <= 1e-06 | 20 | 0.66 | 0.0805 | 0.608<br>(0.57, 0.646) | 1.43<br>(1.26, 1.61) | 3.4<br>(1.37,7.51) |

Notes: Cancer traits are sorted by observed case counts in UKB. \* PRS were scaled to mean = 0 and sd = 1; \*\* Top 1% versus rest

Our observed variations between these cancer PRS likely recapitulates the different genetic architectures of cancers in combination with their prevalences in the discovery and evaluation cohorts. First, the prevalence impacted the ability to identify true associations in the discovery GWAS and also affected our capacity to observe significant effects in the PRS performance evaluation.

#### *Comparison of* Performance across *C*ohorts

The two evaluation cohorts, MGI and UKB, varied in, among other things, their sample sizes, their use of diagnosis code systems, and their recruitment mechanisms, with UKB representing a population-based cohort and MGI an EHR-based, cancer-enriched cohort. We limited a comparison of the cancer PRS to the top-ranked PRS for 19 cancers that were present for both cohorts. We selected the top PRS for each cancer within each cohort, i.e., their GWAS source and method might be different between MGI and UKB.

We noticed the same ranking of AUC values for most cancer PRS but found significantly higher estimates for cancer of brain, cancer of brain / nervous system, colorectal cancer, and prostate cancer in UKB than in MGI (**Figure S4**). The former two estimates might reflect the different underlying GWAS sources, while the latter two might be inflated in UKB due to overlapping samples between their discovery GWAS meta-analyses and the UKB cohort [12, 15]. The other 15 cancers showed a similar ranking of AUC estimates in both cohorts that ranged between “Cancer of other lymphoid, histiocytic tissue” (AUCMGI: 0.527, AUCUKB: 0.532) and highest for “chronic lymphoid leukemia” (AUCMGI: 0.682, AUCUKB: 0.690). AUC values tended to be slightly higher for UKB than for MGI, while confidence intervals were mostly smaller in UKB corresponding to their (often) larger observed effective sample sizes.

A similar comparison of the enrichment of cases in the top 10% versus bottom 90% revealed a lack of power for many cancer PRS in MGI with OR <= 1.3, but a relatively consistent ranking from PRS for pancreatic cancer (MGI OR_Top10%_: 1.37 and UKB OR_Top10%_: 1.64) to prostate cancer (MGI OR_Top10%_: 3.65 and UKB OR_Top10%_: 3.97). Overall enrichment effects were often stronger in UKB compared to MGI, reflecting the larger sample sizes of these cancers but also indicated a disparity between population- and hospital-based controls (**Table 1 & Table S6; Figure S5**). However, when comparing the enrichment of cases for two PRS that were well-powered in both cohorts (PRS for breast cancer and chronic lymphoid leukemia), we found it to be strikingly comparable across all tested percentiles (1, 2, 5, 10, and 25% versus rest; **Figure 2**).

**Figure 2.**
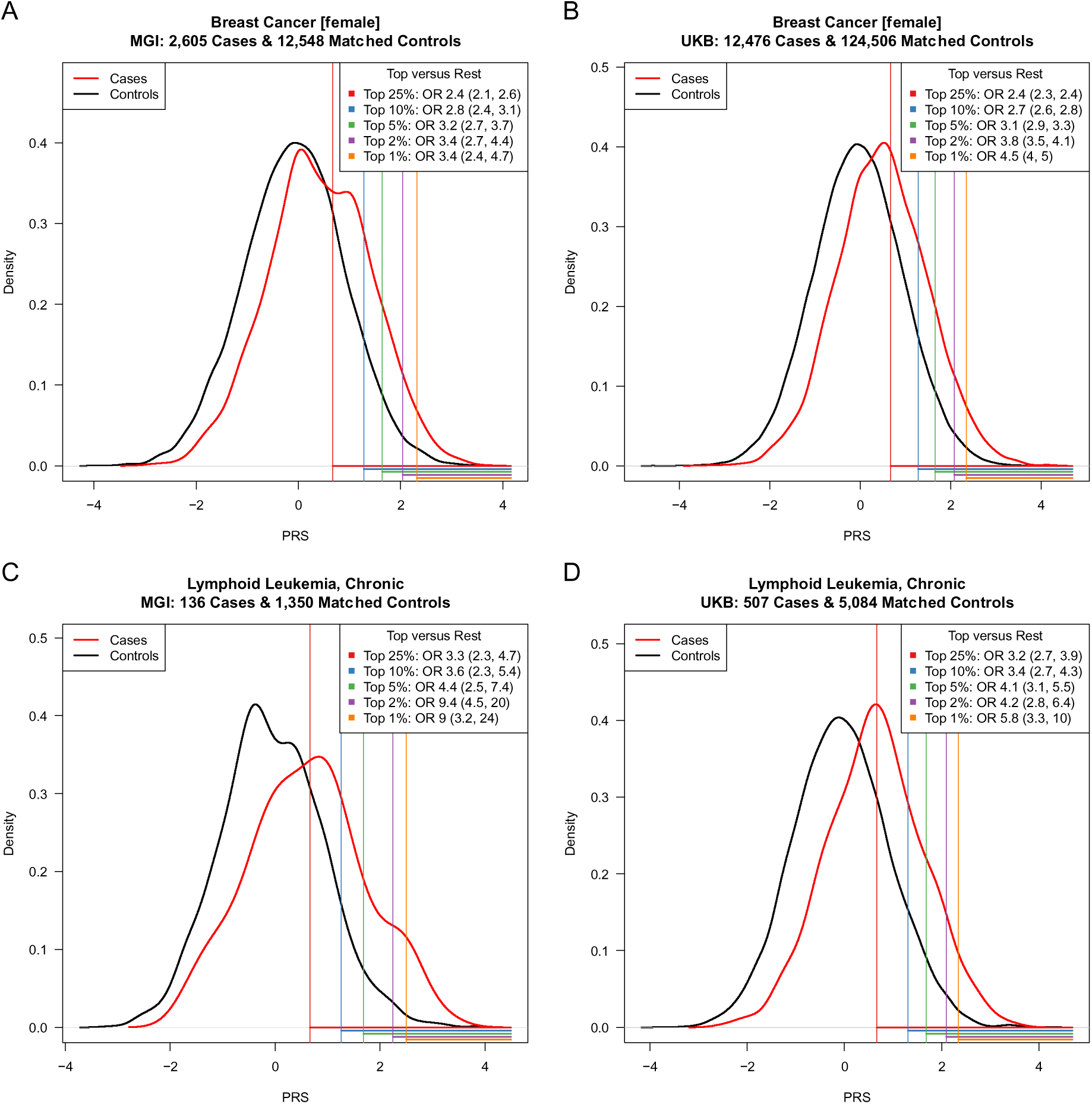
Distribution of breast cancer PRS (A, B), and chronic lymphoid leukemia (C, D) in matched case controls samples in MGI (A, C) and UKB (B, D). Enrichment of cases in five top PRS percentiles is indicated by the shaded areas under the density curves while corresponding Odds ratios (OR) are given in the top right corner of each plot. PRS were standardized.

### Phenome-wide Association Analyses

Beyond case enrichment and risk stratification, PRS can also be used in phenome-wide screenings to uncover secondary trait associations through shared genetic risk factors [31, 32]. These secondary traits might uncover features in the EHR that occur before cancer diagnosis and thus could represent important predictors for cancer outcomes. From the generated PRS for 49 cancer traits, we selected 13 PRS in MGI and 18 PRS in UKB (whose association with their corresponding cancer traits reached phenome-wide significance) for phenome-wide screens of PRS associations. In total, we observed phenome-wide significant associations between 21 cancer PRS and 150 different secondary traits (**Table S5**). We performed “Exclusion-PRS-PheWAS” (i.e., removed primary cancer cases and repeated the phenome-wide analysis) to assess if the identified secondary associations were mainly driven by the primary cancer trait, e.g., through intensified screening or represent post-treatment effects [31]. While the exclusion of cases markedly decreased case counts of secondary traits, we still identified secondary traits that remained significantly associated with the corresponding cancer PRS (**Table 7&8, Table S5**). Most of the secondary traits in MGI that remained phenome-wide significant in the Exclusion-PRS-PheWAS, e.g., skin cancer PRS associated with actinic keratosis or thyroid cancer PRS associated with hypothyroidism, were reported in our previous studies [32, 53]. Due to the larger sample sizes for most traits in UKB compared to MGI (**Table S6**), we observed more and stronger secondary trait associations in UKB PRS-PheWAS. Several secondary trait associations were seen in both cohorts (e.g., hypothyroidism associated with thyroid cancer PRS after exclusion thyroid cancer cases: OR_MGI_ = 0.864 [0.838, 0.89] and OR_UKB_ = 0.896 [0.881,0.912]; **Figure 3 A & C**). We also observed several secondary trait associations exclusively in UKB. Some of these associations, e.g., hyperplasia of prostate associated with cancer of prostate PRS (Exclusion-PRS-PheWAS in UKB: OR 1.07 [95% CI: 1.05, 1.09], P = 2.16E-10), represent known risk factors or presentation features of primary cancers [54, 55]. However, we also observed traits where the cancer relevance was less clear, e.g., varicose veins associated with breast cancer PRS (Exclusion-PRS-PheWAS in UKB: OR 1.05 [95% CI: 1.03,1.07], P = 2.88E-07) (**Figure 3 C & D**; **Table S5)**. Deeper explorations and replications are needed to understand these observed associations and to distinguish between spurious and genuine associations.

**Figure 3.**
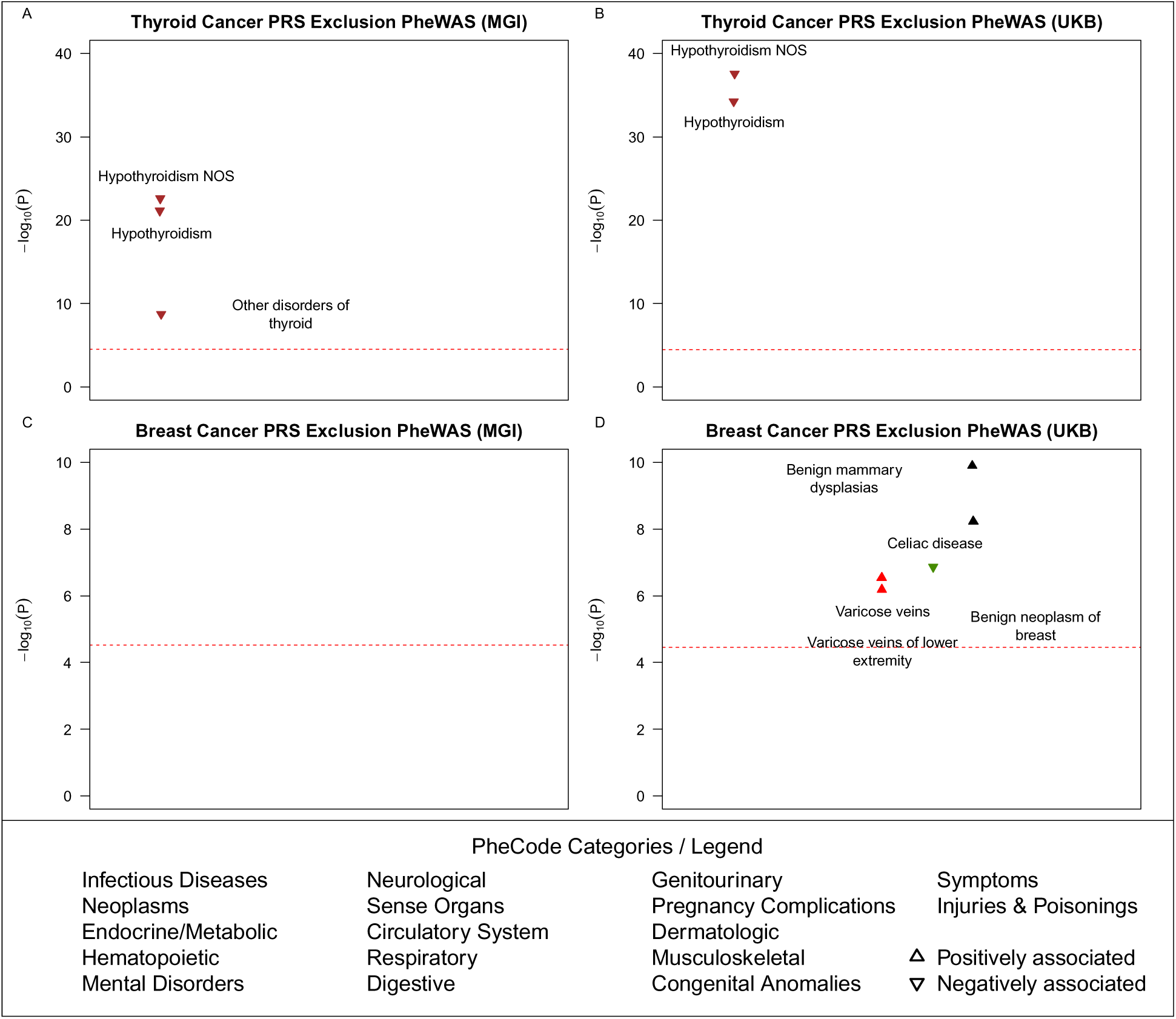
Exclusion PRS-PheWAS in the MGI and UKB phenomes. The horizontal line indicates phenome-wide significance. Only the strongest and phenome-wide significantly associated traits within a category are labelled. Directional triangles indicate whether a phenome-wide significant trait was positively (pointing up) or negatively (pointing down) associated with the PRS.

**Table 7.** Secondary traits associated with primary cancer PRS in MGI. Trait / PRS associations that reached phenome-wide significance (P <= 0.05/1679) in the PRS PheWAS in MGI after excluding the primary and related cancer traits from the analysis are shown

| Cancer trait<br>underlying PRS<br>(PheWAS Code) | Secondary trait<br>associated with PRS<br>(PheWAS Code) | Primary analysis |  | After exclusion* |  |
| --- | --- | --- | --- | --- | --- |
|  |  | OR (95% CI) | P | OR (95% CI) | P |
| Colorectal cancer<br>(153) | Benign neoplasm of<br>colon (208) | 1.1<br>(1.07,1.13) | 2.75E-13 | 1.08<br>(1.05,1.11) | 5.19E-09 |
| Non-melanoma skin<br>cancer (172.2) | Benign neoplasm of<br>skin (216) | 1.07<br>(1.04,1.09) | 1.49E-07 | 0.937<br>(0.91,0.966) | 2.72E-05 |
| Squamous cell<br>carcinoma (172.22) | Actinic keratosis<br>(702.1) | 1.38<br>(1.34,1.44) | 4.54E-68 | 1.24<br>(1.17,1.31) | 1.99E-14 |
| Carcinoma in situ of<br>skin (172.3) | Actinic keratosis<br>(702.1) | 1.29<br>(1.25,1.34) | 6.78E-49 | 1.18<br>(1.12,1.24) | 6.78E-10 |
| Thyroid cancer<br>(193) | Hypothyroidism<br>(244) | 0.926<br>(0.901,0.953) | 1.05E-07 | 0.864<br>(0.838,0.89) | 7.40E-22 |
| Thyroid cancer<br>(193) | Hypothyroidism NOS<br>(244.4) | 0.914<br>(0.887,0.941) | 1.26E-09 | 0.855<br>(0.829,0.882) | 2.46E-23 |
| Thyroid cancer<br>(193) | Other disorders of<br>thyroid (246) | 0.947<br>(0.918,0.978) | 0.000741 | 0.904<br>(0.875,0.934) | 1.88E-09 |

\* primary and related cancer traits excluded from analysis; NOS: not otherwise specified

**Table 8.** Secondary traits associated with primary cancer PRS in UKB. PRS / trait associations that reached phenome-wide significance (P <= 0.05/1419) in the PRS PheWAS in UKB after excluding the primary and related cancer traits from the analysis are show.

| **Cancer trait underlying PRS (PheWAS Code)** | **Secondary trait associated with PRS (PheWAS Code)** |
| --- | --- |
| Colorectal cancer (153) | Neoplasm of unspecified nature of digestive system (158); Malignant neoplasm of other and ill-defined sites within the digestive organs and peritoneum (159); Cancer, suspected or other (195); Malignant neoplasm, other (195.1); Benign neoplasm of colon (208); Diverticulosis and diverticulitis (562); Diverticulosis (562.1); Functional digestive disorders (564); Personal history of diseases of digestive system (564.9); Anal and rectal conditions (565); Anal and rectal polyp (565.1) |
| Pancreatic cancer (157) | Phlebitis and thrombophlebitis (451); Phlebitis and thrombophlebitis of lower extremities (451.2) |
| Cancer of bronchus / lung (165.1) | Cancer of urinary organs (incl. kidney and bladder) (189); Peripheral vascular disease (443); Other specified peripheral vascular diseases (443.8); Peripheral vascular disease, unspecified (443.9); Celiac disease (557.1) |
| Melanomas of skin (172.1) | Benign neoplasm of skin (216); Angina pectoris (411.3); Disorder of skin and subcutaneous tissue NOS (689); Degenerative skin conditions and other dermatoses (702); Actinic keratosis (702.1) |
| Breast cancer [female] (174.1) | Varicose veins (454); Varicose veins of lower extremity (454.1); Celiac disease (557.1); Benign mammary dysplasias (610); Benign neoplasm of breast (610.4) |
| Cancer of uterus (182) | Breast cancer (174); Breast cancer [female] (174.1); Malignant neoplasm of female breast (174.11); Coronary atherosclerosis (411.4); Osteoporosis, osteopenia and pathological fracture (743); Osteoporosis (743.1); Osteoporosis NOS (743.11) |
| Cancer of ovary (184.11) | Phlebitis and thrombophlebitis (451); Celiac disease (557.1); Inflammatory diseases of female pelvic organs (614); Inflammatory disease of cervix, vagina, and vulva (614.5) |
| Cancer of prostate (185) | Celiac disease (557.1); Hyperplasia of prostate (600); Other disorders of prostate (602); Other abnormal blood chemistry (790.6) |
| Cancer of brain and nervous system (191.1) | Benign neoplasm of uterus (218); Uterine leiomyoma (218.1); Cataract (366); Coronary atherosclerosis (411.4); Polyp of female genital organs (622); Polyp of corpus uteri (622.1) |
| Cancer of brain (191.11) | Benign neoplasm of uterus (218); Uterine leiomyoma (218.1); Nontoxic nodular goiter (241); Glaucoma (365); Cataract (366); Ischemic Heart Disease (411); Myocardial infarction (411.2); Angina pectoris (411.3); Coronary atherosclerosis (411.4); Other chronic ischemic heart disease, unspecified (411.8); Excessive or frequent menstruation (626.12) |
| Thyroid cancer (193) | Hypothyroidism (244); Hypothyroidism NOS (244.4) |
| Hodgkin's disease (201) | Diabetes mellitus (250); Type 1 diabetes (250.1); Type 1 diabetes with ketoacidosis (250.11); Type 1 diabetes with ophthalmic manifestations (250.13); Multiple sclerosis (335); Nasal polyps (471); Celiac disease (557.1); Rheumatoid arthritis and other inflammatory polyarthropathies (714); Rheumatoid arthritis (714.1) |
| Cancer of other lymphoid, histiocytic tissue (202) | Thyrotoxicosis with or without goiter (242); Disorders of protein plasma/amino-acid transport and metabolism (270); Disorders of plasma protein metabolism (270.3); Celiac disease (557.1) |
| Non-Hodgkins lymphoma (202.2) | Multiple sclerosis (335); Nasal polyps (471); Celiac disease (557.1); Sarcoidosis (697); Rheumatoid arthritis and other inflammatory polyarthropathies (714); Rheumatoid arthritis (714.1); Ankylosing spondylitis (715.2) |
| Lymphoid leukemia, chronic (204.12) | Celiac disease (557.1) |
| Multiple myeloma (204.4) | Skin cancer (172); Other non-epithelial cancer of skin (172.2) |

### Online Visual Catalog: *Cancer PRSweb* and R package Rprs

In our current study, we compared three PRS construction methods for 68 cancer traits using over 232 sets of GWAS summary statistics. By doing so, we created a large number of PRS in which predictive or enrichment properties differed between GWAS source, PRS method, and/or evaluation cohort. After assessing 1,292 constructed PRS, we found PRS for 49 different cancer traits that we deemed to have predictive value. In our explorations, we established that it could be beneficial to select PRS with certain predictive properties for a specific application instead of using one PRS for all applications. Also, it could be computationally more convenient to use a slightly less powerful PRS based on a fewer number of SNPs than to use a PRS that is based on a few hundred thousand variants. To allow the user the option to explore various PRS constructs, we created PRSweb (http://prsweb.sph.umich.edu), an interactive and intuitive web interface, to explore the available PRS constructs for 49 different cancer traits as well as their performance metrics and suitability for risk stratification, association studies, or other PRS applications.

After an initial selection menu for cancer trait and evaluation cohort (MGI or UKB) PRSweb provides tabularized information about all available PRS, their evaluation metrics (performance, discrimination, calibration, and accuracy) and case enrichment capabilities in five top percentiles of their distributions. The tables, similar to **Tables 3 & 4**, can be sorted, filtered, or downloaded in full. These tables contain detailed information about the underlying GWAS source(s), LD reference panels and are directly linked to downloadable PRS constructs. The PRS construct files include headers with information about the PRS construction (source, version, method, and references) and lists its underlying risk variants, their physical positions, effect/non-effect alleles (forward strand orientation for a given genome assembly), and its weights. Together with the “Rprs” R package (https://github.com/statgen/Rprs) we developed, the construct file will enable the reproduction of PRS association in MGI or UKB and allow a straightforward generation of comparable PRS in external datasets using imputed dosage data in VCF or BCF format.

For phenome-wide predictive PRS (association P_PrimaryCancer_ ≤ 0.05 / [# phenotypes in phenome]), PRSweb also links to PRS-PheWAS results for their evaluation cohort. The PRS-PheWAS result page includes interactive Manhattan plots for PRS-PheWAS and Exclusion-PRS-PheWAS with mouseover information for each tested association. The PheWAS plots can be exported as scalable vector graphics (SVG) and are accompanied by interactive and downloadable result tables that provide PheWAS summary statistics plus sample counts per analyzed phenotype.

We also implemented a search interface for each phenotype/PheCode to provide insights into the ICD-codes underlying the primary cancers as well as the traits of our EHR-derived MGI and UKB phenomes. A methods section describes the applied approaches.

## Discussion

In our study, we constructed and evaluated a large set of cancer PRS using more than 200 different sources of GWAS summary statistics. We applied three common PRS construction methods: GWAS hits, LD pruning/P-value thresholding, and lassosum. While doing so, we created an online repository called “PRSweb” (http://prsweb.sph.umich.edu/) with over 600 PRS for 49 cancer traits.

We observed that construction and resulting performance of PRS depend on multiple factors, including GWAS source, PRS method, and evaluation cohort. Researchers who plan to apply PRS in their projects are often faced with an agony of choice from a set of PRS in the current literature or might not find predictive PRS at all. Furthermore, if PRS are available, a direct comparison of multiple constructs is often not feasible, as their performance can be cohort-specific and limited by available sample size.

To alleviate this situation, we generated “PRSweb” that could serve as a central hub for standardized PRS. PRSweb so far offered a selection and exploration of PRS based on publicly available cancer GWAS data. The platform integrated the evaluation of the rich EHR data of two independent biobanks, MGI and UKB. In our initial version of PRSweb, we focused on cancer traits because MGI is enriched for cancer.

There are several remaining challenges in developing PRS, and we will discuss the following four: access to independent GWAS summary statistics, mapping of trait definitions between discovery and evaluation cohorts, power limitations, and finally, transferability of PRS across cohorts and ancestries.

### Access to independent GWAS summary statistics

Limited accessibility to full summary statistics for cancer GWAS in the published literature resulted in a lack of PRS constructs for many cancers. By systematically integrating openly available cancer GWAS summary statistics, we can also openly share PRS constructs, some with millions of markers, with the research community. However, there are large cancer GWAS datasets used in the cancer research community that are not yet integrated into PRSweb. For example, a recent study analyzed fourteen different cancer types based on summary-level association statistics from larger cancer GWAS consortia [56]. To our knowledge, only the full summary statistics on breast cancer [11], ovarian cancer [9], and prostate cancer [12] were openly shared. We are confident that future versions of PRSweb will be able to integrate summary statistics from other large GWAS consortia, e.g., on chronic lymphocytic leukemia, glioma, melanoma, esophageal, testicular, oropharyngeal, pancreatic, renal, colorectal, endometrial, or lung cancer, some with substantially larger samples sizes than the GWAS used in our current analysis.

With the tendency to form large consortia and to integrate available biobank data comes another challenge, namely the potential overlap between the discovery and evaluation cohorts and, thus, potential overfitting. For our current study, we used GWAS that are to the best of our knowledge, independent from MGI. Since UKB is a popular and widely used resource, the assumption of independence of large GWAS efforts from UKB, does not always hold true as we have seen for the large colorectal cancer GWAS[15]. In the future, the assessment of independence of GWAS from PRS construction will become more challenging, especially when relying on GWAS databases (e.g., the GWAS Catalog), where the distinction of contributing cohorts might not be obvious from a database entry alone. An alternative solution, especially for consortia joining large GWAS, is leave-one-out meta-analysis where in addition to the full meta-analysis results, a separate set of meta-analysis results will be provided for each contributing cohort so that each resulting leave-one-out meta-analysis can be shared and used for PRS generation in that cohort to avoid overfitting.

We anticipate a more accessible landscape of high-quality full GWAS results in the near future, not only for cancer. First, funding agencies are updating their policy regarding access to GWAS summary statistics of funded projects, e.g., the US National Institutes of Health (NIH), (https://grants.nih.gov/grants/guide/notice-files/NOT-OD-19-023.html). Secondly, biobank studies are growing in numbers and size and, when connected to EHR data, enable GWAS for thousands of traits each [53]. In addition, global efforts are forming that will enable even more powerful phenome x GWAS meta-analyses through collaboration, likely reaching sample sizes that can compete with classical disease-specific consortia [57].

### Mapping of Trait Definitions

One of the premises for PRS utility is the resemblance of the original trait in the discovery GWAS with the trait of the evaluation cohort. For our current study, we relied on EHR-based cohorts and defined cancer via PheWAS codes that are adopted from ICD codes. It is important to bear in mind that we used EHR-based diagnosis data that *per se* were not collected for research. Besides misclassification, EHR-derived phenomes might be prone to selection and recruitment biases that can negatively impact power or result in false-positive associations [53]. ICD codes usually serve administrative and billing purposes and often lack the specificity found in the discovery GWAS. Due to the difference in trait definitions, we often had to fall back to the broad phenotype definition in the EHR cohorts and, by doing so, might have negatively influenced the predictive power for PRS [58]. For example, we only had one definition for ovarian cancer in MGI and UKB (Phecode 184.11 “Malignant neoplasm of ovary”) that was defined by ICD9 codes 183.0 and V10.43 as well as by ICD10:C56 and their sub-codes. In contrast, the large GWAS on ovarian cancer included results for nine more refined cancer subtypes: invasive epithelial, low-grade serous, high-grade serous, serous invasive, endometrioid, epithelial, mucinous, low-grade serous and serous borderline ovarian cancer, and ovarian clear cell cancer. For our PRS generation, we used all nine GWAS as separate sources and tested each resulting PRS against the single PheWAS code 184.11. Consequently, the best performing PRS might represent the combination where the discovery GWAS’s trait specificity and the cohort’s trait composition maximized predictive power.

While we were bound to PheWAS code definitions, future PRS explorations and evaluations of growing EHR data should include more refined cancer phenotypes by integrating cancer registry data and/or natural language processing of clinical notes. Still, the chosen phenotype definitions represent valid and common cancer groupings that are frequently used in clinical and research applications [59].

### Power Limitations

For our project, we used data from MGI, a medical center-based cohort, and UKB, a population-based cohort. Due to MGI’s recruitment mechanism through surgery, the observed case counts in MGI reflected the numbers of adult (18+) patients that underwent a surgical procedure and had at least one corresponding cancer diagnosis in their medical records. The case counts in UKB, a rather healthy subset of the older (40-69) British population [60], might be even lower than the population’s cancer prevalence. We observed an enrichment of many cancers in MGI compared to UKB, especially for rarer cancers like thyroid cancer, but generally registered more cases in the UKB because its cohort is ten times larger (**Table S6**). In addition, MGI’s recruitment through surgical procedures likely resulted in a relative depletion of blood cancers (e.g., leukemia, lymphoma, and myeloma), since affected patients undergo surgery less frequently than somatic cancer patients. As a consequence, we often had sufficient power to evaluate and analyze PRS for these diseases in UKB but not MGI.

One may be interested in defining a combined phenotype of “any cancer” for a composite cancer PRS with a maximal sample size. We defined this phenotype in UKB (with 69,190 cases of any cancer; **Tables S6 & S7, Figures S7 & S8**), performed a GWAS that revealed known risk variants for numerous cancers, and created an “any cancer” PRS using our established methods. The lassosum PRS with a choice for 179 variants performed best amongst the constructs (**Table S8**). However, while defining such a composite phenotype, we have to remember that the endpoint is a heterogeneous mix of various cancers, and the discovery will be driven by the cancers with larger numbers of cases or strong risk effects in the discovery (UKB) and evaluation (MGI) cohort. In the PRS PheWAS in MGI, we saw many related traits associated with the overall PRS. No secondary trait reached phenome-wide significance in the exclusion PRS-PheWAS (**Table S9; Figure S9**). We incorporated this overall PRS constructs in Cancer-PRSweb.

Besides accessible sample sizes, the ability to create predictive PRS depends on the cancers’ “chip heritability,” i.e., the variance explained through polygenic variants of genotyped and imputed datasets. A previous study on six common cancers found that chip heritability estimates can vary substantially for cancers (e.g., estimated heritability for prostate cancer: 27%, breast cancer 12%, and pancreatic cancer 7%) [61]. Thus, indicating that even if the most powerful cancer PRS can be generated, other factors play a bigger role, emphasizing the limitations of PRS for personal risk prediction if used on its own without considering other risk factors [62].

Also, genetic architecture affects the choice of PRS construction methods. A recent study estimated the heritability explained by genome-wide significant variants for 14 common cancers and found a wide variability of explained heritability estimates among the analyzed cancer types. For some cancers like testicular cancer, chronic lymphocytic leukemia, prostate, and breast cancer, GWAS hits could explain a large fraction of the chip heritability, while GWAS hits for other cancers like esophageal, colorectal, endometrial, ovarian or lung cancer explained only moderate to very low fractions [56]. Consequently, approaches that only consider GWAS hits might work better for the former, while less conservative p-value thresholds or genome-wide PRS methods might work better for the latter cancer traits.

### Transferability of PRS across cohorts

In our current study, we constructed and evaluated PRS in individuals of broadly European ancestry. However, we recognize the need to also construct and share PRS for non-European ancestry groups, especially because of the limited transferability of PRS across ancestries and ethnicities [7]. The integration of PRS for non-European individuals into our platform PRSweb so far is hampered by the scarcity of GWAS data for diverse ancestry groups [63], and by the limited diversity in MGI and UKB, both encompassing predominantly European ancestry individuals.

Differences in genotyping and sequencing strategies can also negatively impact comparability between studies. Ideally, genotype data in the discovery GWAS, the LD reference panel, and the evaluation cohort should be comparable in quality, density and LD structure for ultimate compatibility. GWAS usually rely on genotyping arrays that can differ in composition and density of variants. Phasing and imputation methods are constantly improving thanks to growing reference panels and refined methods [64] and are essential in harmonizing genotype data across cohorts. However, the achievable accuracy is dependent on the study’s sample size and variant density. Consequently, a PRS that was constructed from a large and marker-dense GWAS might not be directly transferable to smaller, sparser genotype data.

In our current analysis of two genotype datasets that differed in genotype density and sample size, we found that the tuning parameters of PRS established separately for MGI and UKB were ranked similarly in terms of their resulting predictive performance. This indicated that sharing of PRS constructs might represent a feasible and convenient alternative to computationally expensive PRS methods and evaluations.

### Conclusions

By generating PRS from a large collection of freely available cancer GWAS summary statistics and by evaluating them in two independent biobanks, we created the analytical backbone of PRSweb, an online repository for cancer PRS offering detailed constructs and comparisons. We designed PRSweb for transparency and convenience to democratize PRS research. So far, we included PRS constructs and analyses for 49 different cancer traits that showed promising performance in MGI and/or UKB. We anticipate the inclusion of additional PRS constructs and methods in an upcoming version of PRSweb that also will expand our focus beyond cancers.

Several challenges remain in PRS research in terms of access, power, and transferability. Nevertheless, PRS have proven to be a valuable tool for risk stratification, especially if combined with non-genetic risk factors [65–67]. PRS will likely become more powerful with growing sample sizes, better tools, and more diverse resources.

## Methods

### Evaluation cohorts

#### MGI cohort

Adult (18+) participants were recruited through the Michigan Medicine health system while awaiting diagnostic or interventional procedures either during a preoperative visit prior to the procedure or on the day of the procedure that required anesthesia. In addition to coded biosamples and secure, protected health information, participants understood that all EHR, claims, and national data sources linkable to the participant may be incorporated into the MGI databank. Each participant donated a blood sample for genetic analysis, underwent baseline vital sign testing, and completed a comprehensive history and physical assessment (also see Ethics Statement below). We report results obtained from 38,360 unrelated, genotyped patients of inferred recent European ancestry with available integrated EHR data (∼90 % of all MGI participants were inferred to be of recent European ancestry) [68]. The data used in this study included diagnoses coded with the Ninth and Tenth Revision of the International Statistical Classification of Diseases (ICD9 and ICD10) with clinical modifications (ICD9-CM and ICD10-CM), sex, precomputed principal components (PCs), genotyping batch, and age. Data were collected according to the Declaration of Helsinki principles [69]. MGI study participants’ consent forms and protocols were reviewed and approved by the University of Michigan Medical School Institutional Review Board (IRB ID HUM00099605 and HUM00155849). Opt-in written informed consent was obtained.

#### UK Biobank cohort (UKB)

UKB is a population-based cohort collected from multiple sites across the United Kingdom and includes over 500,000 participants aged between 40 and 69 years when recruited in 2006–2010 [16]. The open-access UK Biobank data used in this study included genotypes, ICD9 and ICD10 codes, inferred sex, inferred White British ancestry, kinship estimates down to third degree, birthyear, genotype array, and precomputed principal components of the genotypes. **Table 1** provides some descriptive statistics of the MGI and UK Biobank samples.

### Genotyping, sample quality control and imputation

#### MGI

DNA from 47,364 blood samples was genotyped on customized Illumina Infinium CoreExome-24 bead arrays and subjected to various quality control filters, resulting in a set of 392,323 polymorphic variants. Principal components and ancestry were estimated by projecting all genotyped samples into the space of the principal components of the Human Genome Diversity Project reference panel using PLINK (938 individuals) [70, 71]. Pairwise kinship was assessed with the software KING [72], and the software fastindep was used to reduce the data to a maximal subset that contained no pairs of individuals with 3rd-or closer degree relationship [73]. We removed participants without EHR data and participants not of recent European descent from the analysis, resulting in a final sample of 38,360 unrelated subjects. Additional genotypes were obtained using the Haplotype Reference Consortium reference panel of the Michigan Imputation Server [58] and included over 24 million imputed variants with R^2^ ≥0.3 and minor allele frequency (MAF) ≥0.01%. Genotyping, quality control, and imputation are described in detail elsewhere [68].

#### UK Biobank

We used the UK BioBank Imputed Dataset (v3, https://www.ebi.ac.uk/ega/datasets/EGAD00010001474) and limited analyses to the documented 408,961 White British [74] individuals and 47,836,001 variants with imputation information score >= 0.3 and MAF >= 0.01% of which 22,846,729 overlapped with the imputed MGI data (see above). Two random subsets of 5,000 and 10,000 unrelated, White British individuals were used for LD analyses of UKB-based summary statistics.

### Phenome generation

#### MGI

The MGI phenome was used as the discovery dataset and was based on ICD9-CM and ICD10-CM code data for 38,360 unrelated, genotyped individuals of recent European ancestry. Longitudinal time-stamped diagnoses were recoded to indicators for whether a patient ever had given a diagnosis code recorded by Michigan Medicine. These ICD9-CM and ICD10-CM codes were aggregated to form up to 1,857 PheWAS traits using the PheWAS R package (as described in detail elsewhere [68, 75]). For each trait, we identified case and control samples. To minimize differences in age and sex distributions, avoid extreme case-control ratios, and reduce the computational burden, we matched up to 10 controls to each case using the R package “MatchIt” [76]. Nearest neighbor matching was applied for age and the first four principal components of the genotype data (PC1-4) using Mahalanobis distance with a caliper/width of 0.25 standard deviations. Exact matching was applied for sex and genotyping array. A total of 1,689 case-control studies with >50 cases were used for our analyses of the MGI phenome.

#### UK Biobank

The UK Biobank phenome was used as a replication dataset and was based on ICD9 and ICD10 code data of 408,961 White British [74], genotyped individuals that were similarly aggregated to PheWAS traits as MGI (as described elsewhere [77]). In contrast to MGI, there were many pairwise relationships reported for UKB participants.

To retain a larger effective sample size for each phenotype, we first selected a maximal set of unrelated cases for each phenotype (defined as no pairwise relationship of 3^rd^ degree or closer [11, 73]) before selecting a maximal set of unrelated controls unrelated to these cases. Similar to MGI, we matched up to 10 controls to each case using the R package “MatchIt” [76]. Nearest neighbor matching was applied for birthyear and PC1-4 (Mahalanobis-metric matching; matching window caliper/width of 0.25 standard deviations), and exact matching was applied for sex and genotyping array. A total of 1,419 case-control studies with >50 cases each were used for our analyses of the UK Biobank phenome.

On average, we were able to match 9 controls per case in the MGI phenome and 9.9 controls per case in the UKB phenome. Additional phenotype information for MGI and UK Biobank is included in (**Table S6**).

### PRS Structure

PRS combine information across a defined set of genetic loci, incorporating each locus’s association with the target trait. The PRS for patient j takes the form PRS*_j_*=∑*_i_β_i_G_ij_* where *i* indexes the included loci for that trait, weight *βi* is the log odds ratios retrieved from the external GWAS summary statistics for locus *i,* and *G_ij_* is a continuous version of the measured dosage data for the risk allele on locus *i* in subject *j.* In order to construct a PRS, one must determine which genetic loci to include in the PRS and their relative weights. Below, we obtain GWAS summary statistics from several different sources, resulting in several sets of weights for each trait of interest. For each set of weights, we consider several strategies for determining which genetic loci to include in the PRS construction.

### Sources of GWAS summary statistics

For each of 68 cancers of interest, we collected GWAS summary statistics from up to three different sources: (1) merged genome-wide significant association signals published in the NHGRI EBI GWAS Catalog [78] if available; (2) large cancer GWAS meta-analysis if available; and (3) publicly available GWAS summary statistics of phenome x genome screening efforts of the UK Biobank data [77] (see **Web Resources; Figure 1**). If needed, we used LiftOver to convert coordinates of GWAS summary statistics to human genome assembly GRCh37 (https://genome-store.ucsc.edu/).

#### GWAS Catalog

We downloaded previously reported GWAS variants from the NHGRI-EBI GWAS Catalog (file version: r2019-05-03, https://www.ebi.ac.uk/gwas/) [78, 79]. Single nucleotide polymorphism (SNP) positions were converted to GRCh37 using variant IDs from dbSNP (build 151; UCSC Genome Browser, http://genome.ucsc.edu/) after updating outdated dbSNP IDs to their merged dbSNP IDs.

Entries with missing risk alleles, risk allele frequencies, or SNP-disease odds ratios were excluded. If a reported risk allele did not match any of the reported forward strand alleles of a non-ambiguous SNP (not A/T or C/G) in the imputed MGI genotype data (which correspond to the alleles of the imputation reference panel), we assumed minus-strand designation and corrected the effect allele to its complementary base of the forward strand. Entries with a reported risk allele that did not match any of the alleles of an ambiguous SNP (A/T and C/G) in our data were excluded at this step. We only included entries with broad European ancestry (as reported by the NHGRI-EBI GWAS Catalog) to match ancestries of discovery GWAS and target cohorts (MGI and UKB). As a quality control check, we compared the GWAS Catalog reported risk allele frequencies (RAF) with the RAF in MGI individuals. We then excluded entries whose RAF deviated more than 15%. This chosen threshold is subjective and was based on clear differentiation between correct and likely flipped alleles on the two diagonals (**Figure S6**), as noted frequently in GWAS meta-analyses quality control procedures [80]. For SNPs with multiple entries, we kept the SNP with the most recent publication date (and smaller p-value, if necessary) and excluded the others.

#### Large GWAS meta-analyses

We downloaded full GWAS summary statistics made available by the “Breast Cancer Association Consortium” (BCAC) [11], the “Prostate Cancer Association Group to Investigate Cancer Associated Alterations in the Genome” (PRACTICAL) [12], and the “Ovarian Cancer Association Consortium” (OCAC) [9]. In addition, we extracted partial GWAS summary statistics that accompanied recent publications but were incomplete, i.e. reporting only SNPs below a certain p-value threshold [15, 81–83]. GWAS summary statistics were harmonized and, if needed, lifted over to human genome assembly GRCh37. In this paper, this source is referred to as “Large GWAS”.

#### UK-Biobank-based GWAS

We downloaded UK Biobank based GWAS summary statistics from two public repositories.

The first set of UK Biobank GWAS summary statistics were based on the analysis of up to 408,961 White British European-ancestry samples (https://www.leelabsg.org/resources). SNP-disease odds ratios were estimated using logistic mixed modeling adjusting for sample relatedness, and p-values were estimated using saddlepoint approximations (SAIGE method [17]) to calibrate the distribution of score test statistics and, thus, control for unbalanced case-control ratios. The underlying phenotypes were auto-curated phenotypes based on the PheCodes of the PheWAS R package [68, 75, 77] similar to the phenomes used in our study and in the following are referred to as “UKB PHECODE” (**Table S1**).

The second set of UK Biobank GWAS summary statistics were based on a linear regression model of up to 361,194 unrelated White British samples adjusting for relevant covariates (https://github.com/Nealelab/UK_Biobank_GWAS). Three phenotype models were used in their analyses: (1) “PHESANT”: auto-curated phenotypes using PHEnome Scan ANalysis Tool (https://github.com/MRCIEU/PHESANT), (2) “ICD10”: individuals with the same ICD10 category code (first three characters, e.g. “C50”) were used as cases while all non-coded individuals were treated as controls, and (3) “FINNGEN”: curated phenotypes / endpoints based on definitions of the Finngen consortium (https://www.finngen.fi/en/researchers/clinical-endpoints). In addition to the “UKB PHECODE” (described above), these three latter sources are referred to as “UKB PHESANT”, “UKB ICD10” and “UKB FINNGEN”, respectively (**Table S1**).

### PRS Construction

For each set of GWAS summary statistics from the above-mentioned sources and each cancer, we develop up to seven different PRS using three different construction methods (**Figure 1**). Our goal of this approach was to compare multiple PRS methods and find the method that works best for the various types of GWAS summary statistics.

For the first two construction strategies, we performed LD clumping/pruning of variants with p-values below 10^-4^ by using the imputed allele dosages of 10,000 randomly selected samples and a pairwise correlation cut-off at r^2^ < 0.1 within 1Mb window. Using the resulting loci, we defined up to five sub-sets of variants with p-values below different thresholds (<5×10^-9^ to <5×10^-5^). These were used to construct a PRS tied to each threshold, where the PRS associated with p-values less than 5×10^-8^ is sometimes denoted as “GWAS hits.” For the second PRS construction method, we construct many different PRS across a fine grid of p-value thresholds. The p-value threshold with the highest cross-validated pseudo-R2 (see **PRS Evaluation** below) was used to define the more optimized “Pruning and Thresholding (P & T)” PRS.

As an alternative to the p-value thresholding and “P&T” PRS construction strategies, we also used the software package “lassosum” [22] to define a third type of PRS for GWAS sources with full summary statistics. Lassosum obtains PRS weights by applying elastic net penalization to GWAS summary statistics and incorporating LD information from a reference panel. Here, we used 5,000 randomly selected, unrelated samples as the LD reference panel. We applied a MAF filter of 1 % and, in contrast to the other two approaches, only included autosomal variants that overlap between summary statistics, LD reference panel, and target panel. Each “lassosum” run resulted in up to 76 combinations of the elastic net tuning parameters s and λ, and consequently, in 76 SNP sets with corresponding weights used to construct 76 PRS. We then selected the PRS with the highest cross-validated pseudo-R^2^ to define the “lassosum” PRS.

For each cancer and set of GWAS summary statistics, this approach resulted in up to seven PRS, where PRS with less than 5 included variants were excluded and the available GWAS summary statistics limited the available PRS construction techniques in some cases. Using the R package “Rprs” (https://github.com/statgen/Rprs), the value of each PRS was then calculated for each MGI participant and, if the GWAS source was not based on UKB, also for each UKB participant. For comparability of association effect sizes corresponding to the continuous PRS across cancer traits and PRS construction methods, we centered PRS values in MGI and UKB to their mean and scaled them to have a standard deviation of 1.

### PRS Evaluation

For the PRS evaluations, we fit the following model for each PRS and cancer phenotype without adjusting for covariates:

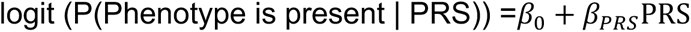

We performed a 5-fold cross validation with the R package “caret” [84] to obtain fitted predictors for the actual PRS evaluations. We used Nagelkerke’s pseudo-R^2^ [85] to select the tuning parameters within the “P&T” and lassosum construction methods (P-value for “P&T” SNP sets; s and λ for lassosum) and kept the PRS with the highest pseudo-R^2^ for further analyses. For each PRS derived for each GWAS source/method combination, we assessed the following performance measures relative to observed disease status in MGI and UKB:

(1) overall performance with Nagelkerke’s pseudo-R^2^ using R packages “rcompanion” [85], (2) accuracy with Brier score using R package “DescTools” [86]; (3) ability to discriminate between cases and controls as measured by the area under the receiver operating characteristic (ROC) curve (denoted AUC) using R package “pROC” [87] and (4) calibration using Hosmer-Lemeshow Goodness of Fit test in the R package “ResourceSelection” [88–90].

### PRS Association Testing

Next, we assessed the strength of the relationship between these PRS and the traits they were designed for. To do this we fit the following model for each PRS and cancer phenotype adjusting for various covariates:

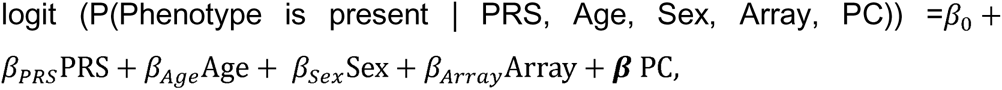

where the PCs were the first four principal components obtained from the principal component analysis of the genotyped GWAS markers, where “Age” was the age at last observed diagnosis in MGI and birthyear in UKB and where “Array” represents the genotyping array. Our primary interest is *β_PRS_*, while the other factors (Age, Sex and PC) were included to address potential residual confounding and do not provide interpretable estimates due to the preceding application of case-control matching. Firth’s bias reduction method was used to resolve the problem of separation in logistic regression (Logistf in R package “EHR”) [91–93].

To study the ability of the PRS to identify high risk patients, we fit the above model but replacing the PRS with an indicator for whether the PRS value was in the top 1, 2, 5, 10, or 25% among the matched case control cohort.

### Phenome-wide Exploration of PRS Associations

We selected PRS that were strongly associated with the cancer trait they were designed for phenome-wide association exploration in the phenomes of MGI and UKB for (p-value ≤ (0.05 / [#phenotypes in corresponding phenome]); see below).

We conducted PheWAS in MGI and also UKB (if the GWAS source was not based on UKB) to identify additional, secondary phenotypes associated with the PRS [31]. To evaluate PRS-phenotype associations, we conducted Firth bias-corrected logistic regression by fitting model of equation 1 above for each PRS and each phenotype of the corresponding phenome. To adjust for multiple testing, we applied the conservative phenome-wide Bonferroni correction according to the total number of analyzed PheWAS codes (MGI: 1,689 phenotypes; UKB: 1,419 phenotypes). In Manhattan plots, we present –log10 (*p*-value) corresponding to tests of *H*_0_: *β_PRS_* = 0. Directional triangles on the PheWAS plot indicate whether a phenome-wide significant trait was positively (pointing up) or negatively (pointing down) associated with the PRS.

To investigate the possibility of the secondary trait associations with PRS being completely driven by the primary trait association, we performed a second set of PheWAS after excluding individuals affected with the primary or related cancer traits for which the PRS was constructed, referred to as “Exclusion-PRS-PheWAS” as described previously [68].

### Online Visual Catalog: *PRSweb*

The online open access visual catalog *PRSweb* was implemented using Grails, a Groovy- and Java-based backend logic, to integrate interactive visualizations and MySQL databases. Interactive PheWAS plots are drawn with the JavaScript library “LocusZoom.js” which is maintained by the UM Center for Statistical Genetics (https://github.com/statgen/locuszoom) and offers dynamic plotting, automatic plot sizing, and label positioning. Additional data-driven visualizations (e.g. temporal relationship plots) were implemented with the JavaScript library “D3.js”.

Unless otherwise stated, analyses were performed using R 3.6.1 [94].

## Data availability

Data cannot be shared publicly due to patient confidentiality. The data underlying the results presented in the study are available from University of Michigan Medical School Central Biorepository at https://research.medicine.umich.edu/our-units/central-biorepository/get-access and from the UK Biobank at http://www.ukbiobank.ac.uk/register-apply/ for researchers who meet the criteria for access to confidential data.

## Acknowledgements

The authors acknowledge the Michigan Genomics Initiative participants, Precision Health at the University of Michigan, the University of Michigan Medical School Data Office for Clinical and Translational Research, the University of Michigan Medical School Central Biorepository, and the University of Michigan Advanced Genomics Core for providing data storage, management, processing, and distribution services, and the Center for Statistical Genetics in the Department of Biostatistics at the School of Public Health for genotype data curation, imputation, and management in support of the research reported in this publication/grant application/presentation. Part of this research has been conducted using the UK Biobank Resource under application number 24460.

This material is based in part upon work supported by the National Institutes of Health/NIH (NCI P30CA046592) and by the National Science Foundation under grant number DMS-1712933. Any opinions, findings, and conclusions or recommendations expressed in this material are those of the author(s) and do not necessarily reflect the views of the National Science Foundation.

## Author contributions

Conceptualization: L.G.F., B.M.; Data Curation: L.G.F.; Formal Analysis: L.G.F., R.B.P.; Funding Acquisition: L.G.F., B.M.; Investigation: L.G.F.; Methodology: L.G.F., L.J.B., B.M.; Project Administration: L.G.F., B.M.; Resources: L.G.F., B.M.; Software: L.G.F., S.P., L.J.B., P.V., M.S., R.B.P., D.T.; Supervision: L.G.F., B.M.; Visualization: L.G.F., S.P., P.V.; Writing – Original Draft Preparation: L.G.F., B.M.; Writing – Review & Editing: L.G.F., S.P., L.J.B., P.V., M.S., R.B.P., D.T., X.Z., B.M.

## Materials & Correspondence

(L.G.F.), (B.M.)

