## Supplementary Figures 1 - 9 for "Cancer PRSweb – an Online Repository with Polygenic Risk Scores (PRS) for Major Cancer Traits and Their Phenome-wide Exploration in Two Independent Biobanks"

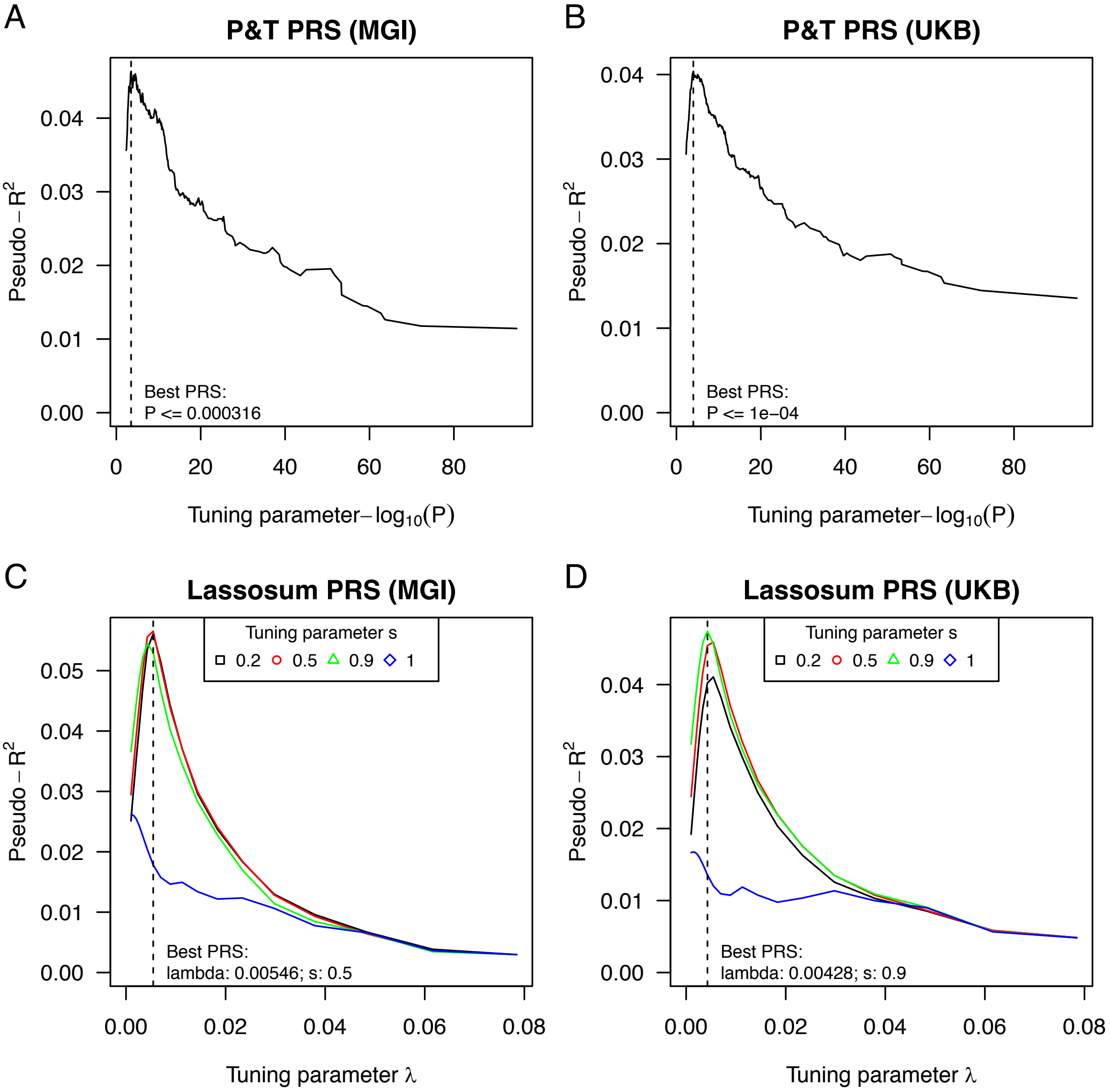


**Figure S1: Tuning Parameter Optimization**: Tuning parameter optimization for the large GWAS based breast cancer PRS with Lassosum (A and B) and "P&T" approach (C & D) for MGI (A & C) and UKB (B & D).

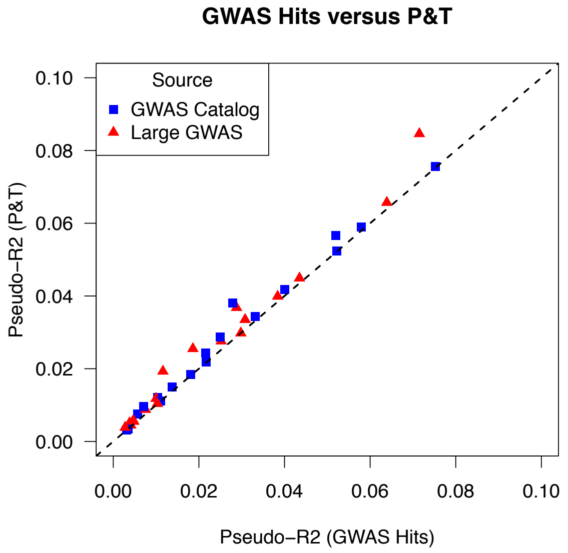


**Figure S2: P&T versus GWAS Catalog hits.** Pairwise comparison of the two PRS methods P&T and "GWAS hits" (P <= 5x10-8) using GWAS Catalog entries as input. 36 PRS for 20 cancer traits (18 MGI PRS and 18 UKB PRS) are shown. Dashed line: identity line.

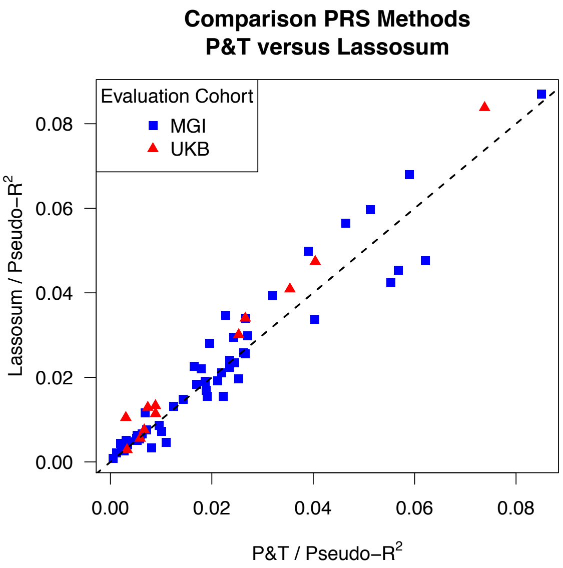


**Figure S3: "P&T versus Lassosum".** Pairwise comparison of the two PRS methods P&T and Lassosum using pseudo-$R^2$. 70 GWAS sources where P&T and Lassosum-based PRS were positively and nominally significant associated with their cancer trait in MGI (blue; 58 PRS) and UKB (red; 12 PRS) are shown. Dashed line: identity line.

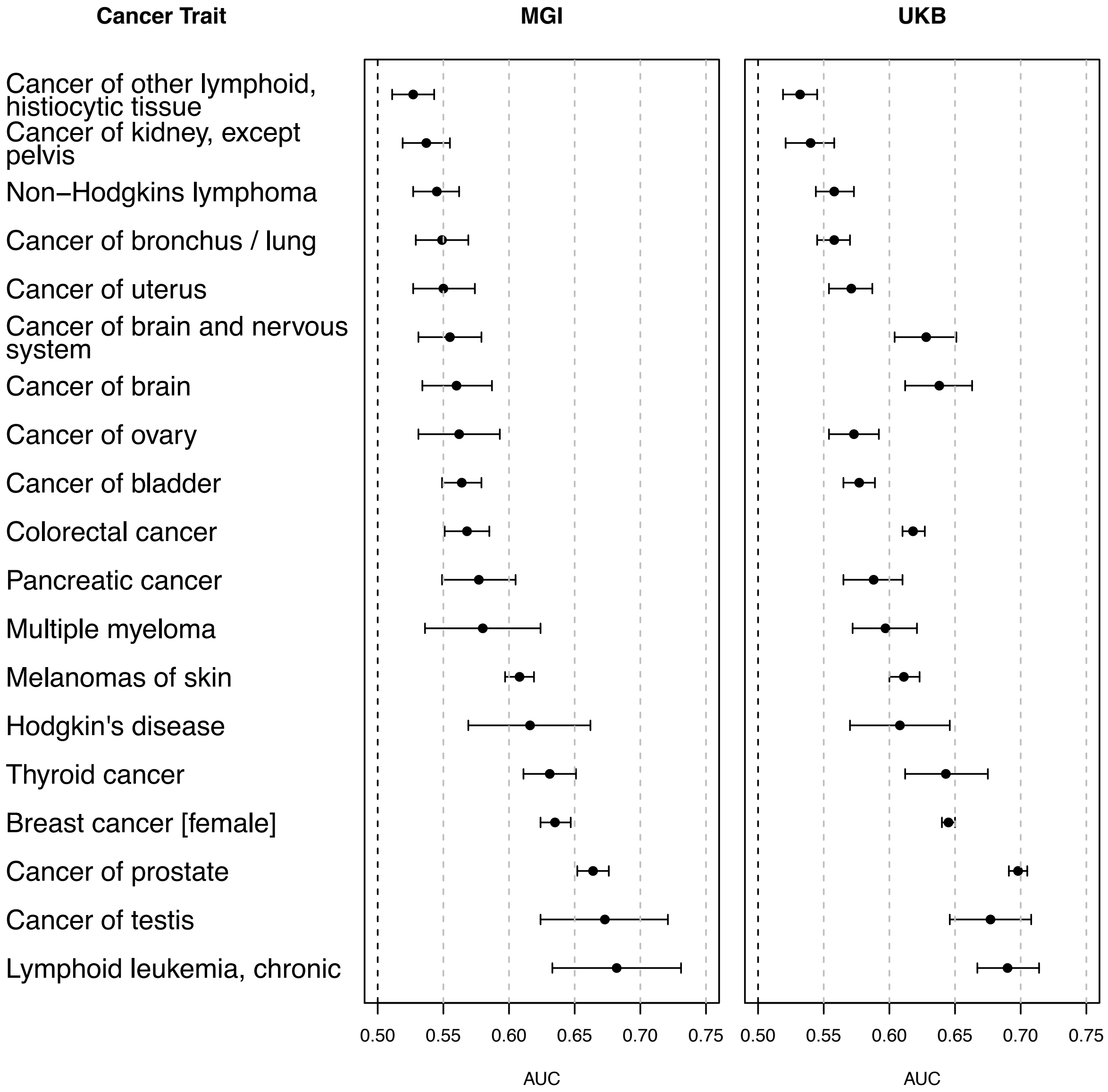


**Figure S4:** AUC of the top ranked PRS for 19 cancers that were present for MGI (left) and UKB (right). AUC values (dots) and their 95% confidence intervals are shown.

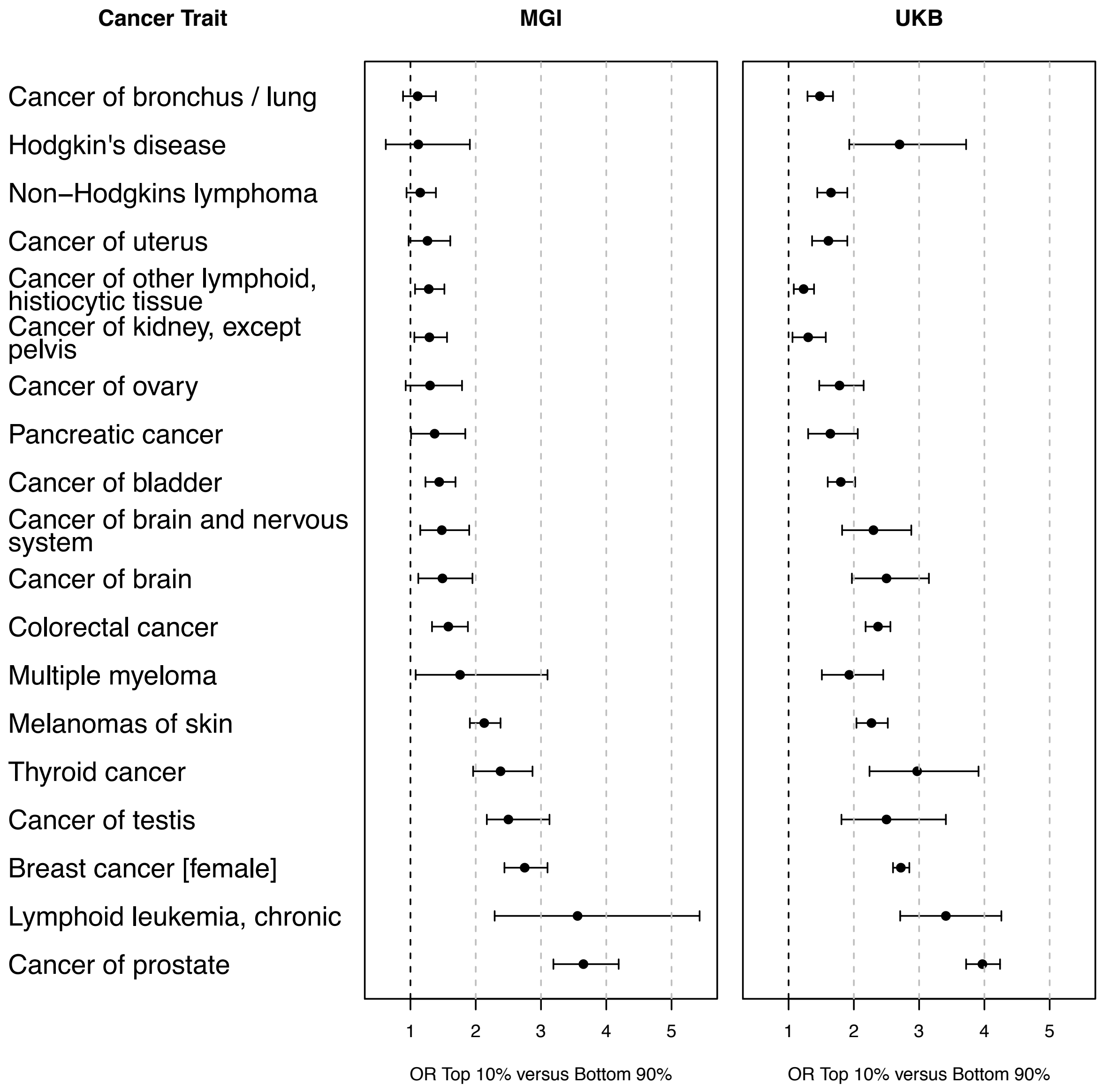


**Figure S5:** Case enrichment of the top ranked PRS for 19 cancers that were present for MGI (left) and UKB (right). Odds ratios (OR, top 10% versus bottom 90% of PRS distribution) and their 95% confidence intervals are shown.

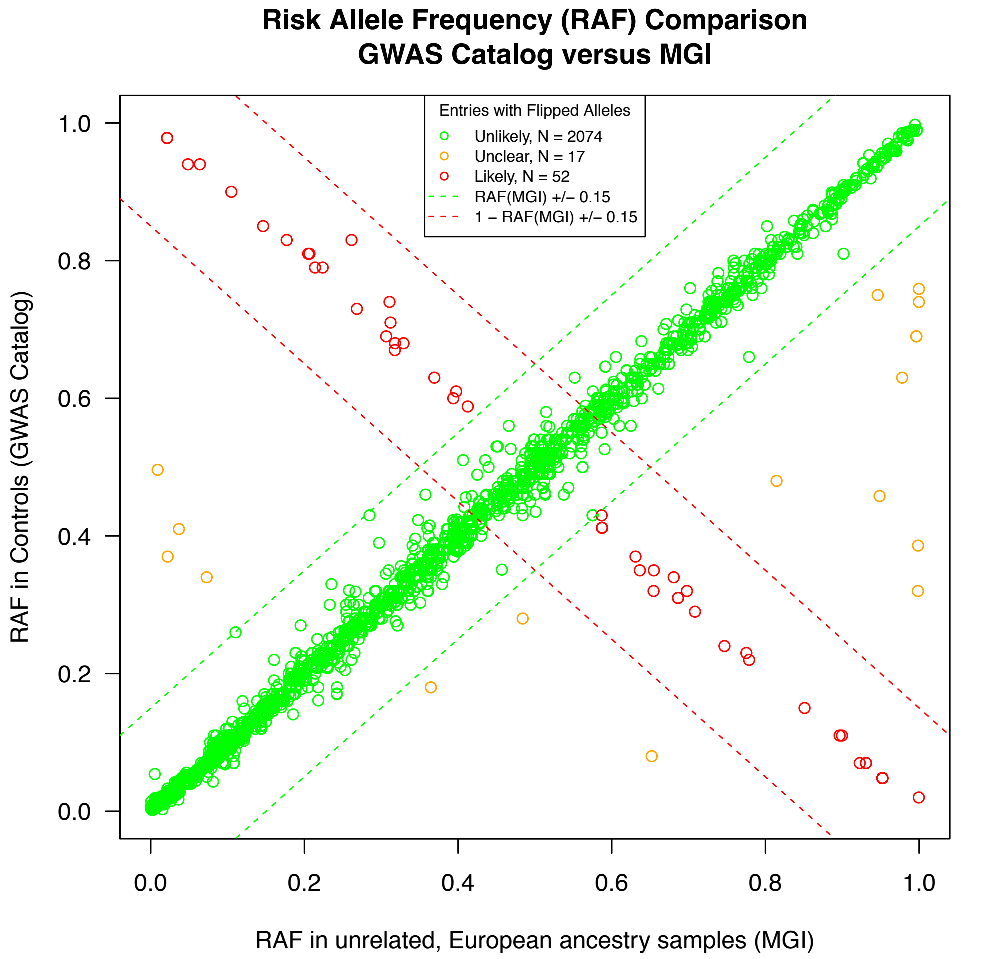


**Figure S6** Comparison of the Risk Allele Frequencies in the GWAS Catalog vs. MGI

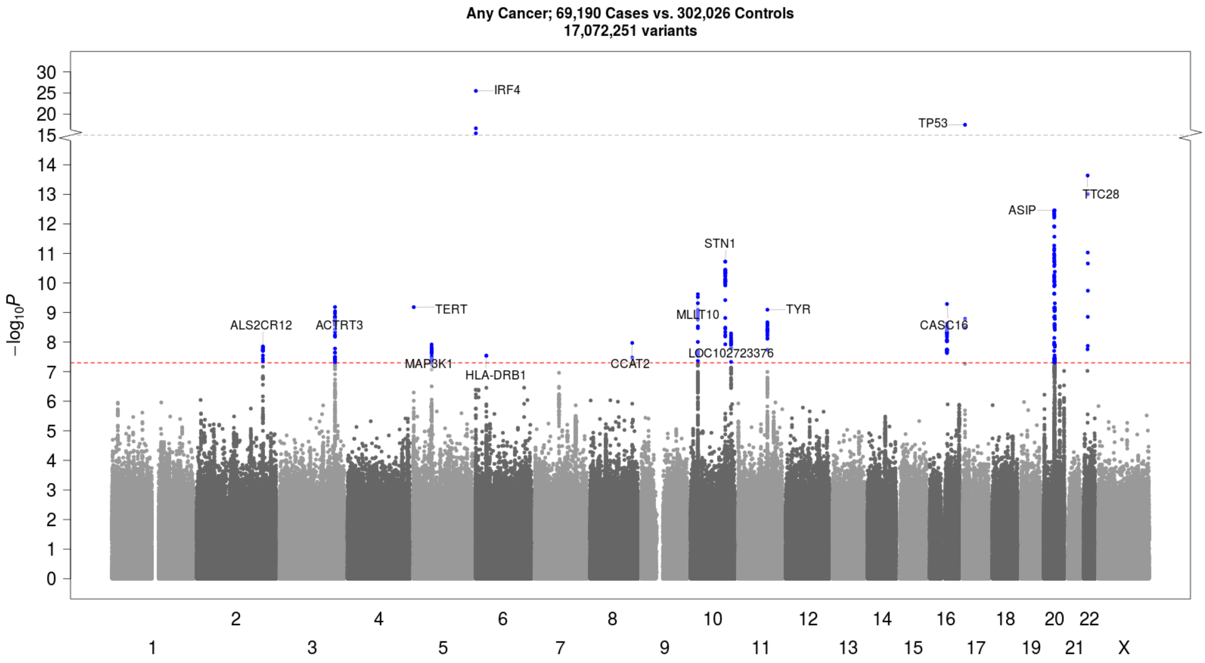


**Figure S7**: Manhattan plot of UKB GWAS on 69,190 cases with any cancer versus 302,026 controls. SNPs with P < 5x10-8 are highlighted in blue. Candidate loci are named after the nearest gene closest to the strongest signal.

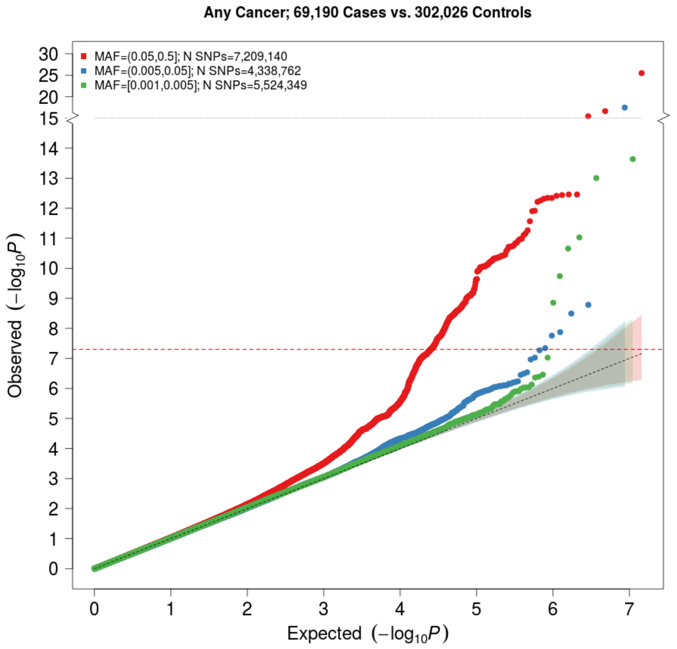


**Figure S8**: QQ plot of UKB GWAS on 69,190 cases with any cancer versus 302,026 controls. -log10(P-values) are stratified by minor allele frequency (MAF) bins.

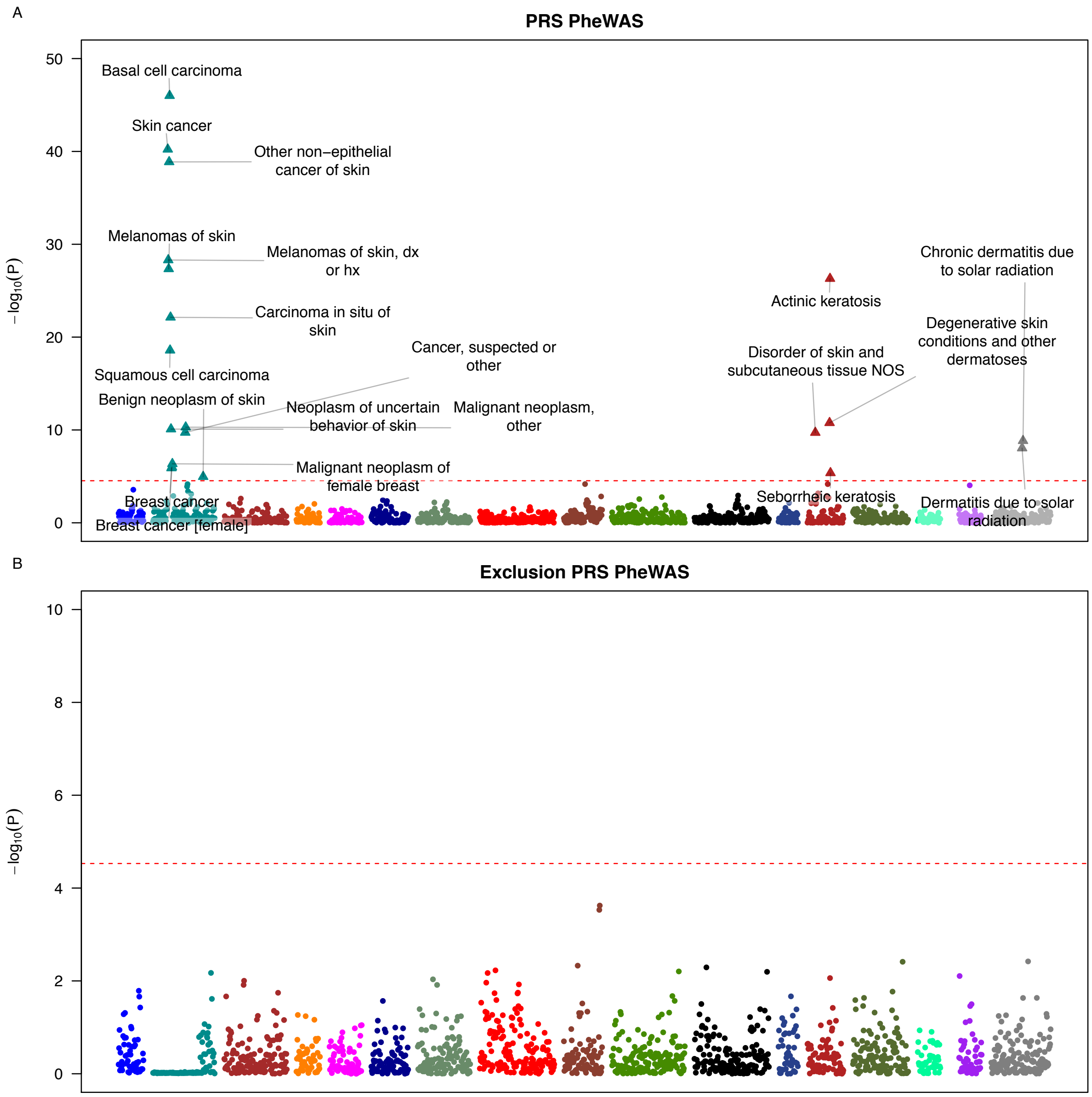


**Figure S9**: PRS PheWAS plot of the 'any cancer' lassosum PRS in MGI before (top) and after (bottom) excluding 20,751 MGI individuals with 'any cancer'.
